# Olfactory learning potentiates long-range cortical GABAergic inputs onto adult-born neurons

**DOI:** 10.64898/2026.05.29.728424

**Authors:** Enzo Peroni, Claire-Hélène De Badts, Gabriel Lepousez, Alya Hiridjee, Justin Renvoise, Ilta Lafosse, Sébastien Wagner, Elise Jacquemet, Pierre-Marie Lledo, Mariana Alonso, Antoine Nissant

## Abstract

Adult neurogenesis in the olfactory bulb (OB) contributes to structural and functional plasticity, influencing olfactory perception, learning, and memory. Adult-born granule cells (abGCs) exhibit unique morphological, electrophysiological, and synaptic properties compared to their neonatally born counterparts, suggesting a specialized role in olfactory processing. In the OB, such processing relies both on sensory inputs from the olfactory epithelium as well as top-down cortical feedback, which encompass both glutamatergic and GABAergic projections from the olfactory cortex back to the OB. While abGCs are known to integrate both bottom-up sensory inputs and top-down cortical projections, the specific connectivity and functional influence of cortical GABAergic inputs on abGCs remain largely unexplored. In this study, we investigated whether activity of cortical GABAergic projections is modulated by olfactory learning, how they impact olfactory behavior and whether these connections selectively influence mature abGCs. Using *in vivo* fiber photometry following odor-reward associative conditioning, we found odor- and reward-dependent activity of cortical GABAergic projections during learning session. Furthermore, their functional role was revealed using optogenetic activation which impaired both the acquisition and the reversal of an odor-reward association. *Ex vivo* patch-clamp recordings demonstrated that olfactory learning potentiates cortical GABAergic inputs specifically onto abGCs, and morphological analysis confirmed that learning increases the number of cortical GABAergic synapses. These findings highlight a novel mechanism by which top-down inhibitory control from the olfactory cortex selectively targets abGC activity during olfactory learning. Our results provide new insights into the functional specialization of abGCs and their role in adaptive olfactory behaviors.

## Material and methods

### Animals

Adult Wild-Type (WT, n=42) C57BL/6J and vGAT::Cre (Slc32a1^tm(cre)Lowl^, MGI ID: 5141270, maintained on a C57BL/6J background, n=120) mice were used in this study. Mice were housed (2-5 per cage), under standard housing conditions (23 ± 1 °C; humidity 40%) in a 14/10h light/dark cycle with dry food and water available ad libitum except during behavioral experiments. All behavioral tests were conducted during the light period (10am – 7pm). All procedures were performed in compliance with the French application of the European Communities Council Directive of 22 September 2010 (2010/63/EEC), approved by the French Ministry of Research (APAFIS #41596-2023031312178181 v2), the local ethics committee (CETEA 89, project dap220049) and were reviewed by the Animal Welfare Committee of Institut Pasteur. We used the minimum number of animals, estimated from our previous knowledge in performing the same type of experiments. Both female and male mice were used in all experiments in similar numbers.

### Viral injections

All viral vectors used here are presented in Table 1. For adult mice injections, P60 (postnatal day 60) mice were anesthetized intraperitoneally with a ketamine (100-150 mg/kg) and xylazine (5 mg/kg) mixture and placed in a stereotaxic apparatus. After local anesthesia and skin incision, a small craniotomy was performed over the targeted region on right and left hemispheres and viral solutions were injected (See Table 2 for stereotaxic coordinates) through a glass micropipette connected to a Nanoject III microinjector (Drummond Scientific). For neonatal mice, P6 (postnatal day 6) pups were anesthetized with isofluorane (3.5%; 372 mL/min; Iso-Vet, Piramal Healthcare) and positioned in a stereotaxic frame using a homemade cast. Small craniotomies were drilled above the injection sites with a needle, and bilateral viral injections (350 nL per site) were made into the RMS (See Table 2 for stereotaxic coordinates). The skin was closed with adhesive cyanoacrylate (Vetbond; 3M). Pups were placed back with their mother after recovery (30–60 min) on a warm pad. After weaning (at P21), mice were segregated by gender. One or two months after lentiviral injection, a second virus was injected in the anterior olfactory cortex (AOC, see Table 2 for stereotaxic coordinates). After all stereotaxic procedures, cranial skin was sutured, and animals were left to recover on a heated pad until complete wakefulness. Paxinos brain atlas was used as reference to verify injection sites in each animal.

**Table 1.**
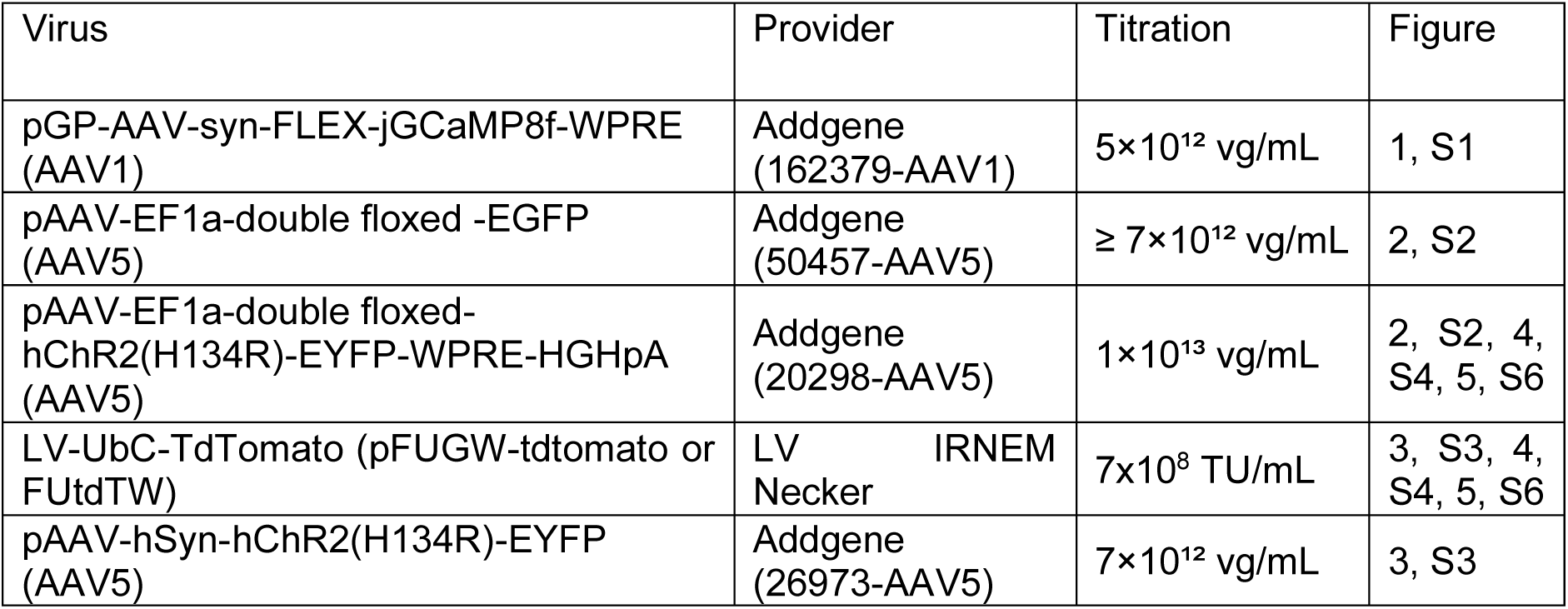
Viral Vectors.

**Table 2.** Sterotaxic coordinates.

| Region | Coordinates | Volume injected | Figure |
| --- | --- | --- | --- |
| RMS | P60<br>AP: +3.3<br>ML: +/-0.82<br>DV: -2.9 from brain surface | P60<br>200nL | 3, S3, 4, S4, 5, S6 |
|  | P6<br>AP: +2.4<br>ML: +/-0.60<br>DV: -2.7 from skull surface | P6<br>350 nL |  |
| AOC | AP: +1.9<br>ML: +/-2.1<br>DV: -3.8/4.0 from brain surface | 200nL per site | 3, S3, 4, S4, 5, S6 |
| AONp | AP: +1.9<br>ML: +/-1.7<br>DV: -4.6 from Bregma | 200nL per site | 1,2 |
| Olfactory peduncle (Fibers implantation) | AP: + 2.9<br>ML: +/- 1.4<br>DV: - 2.5 from brain surface | NA | 1,2 |

### Fiber’s implantation

During stereotaxic procedures, immediately after viral injections, optic fibers (multimode, 430µm diameter, NA 0.5, LC zirconia ferrule, Doric Lenses) were bilaterally implanted above the olfactory peduncle (See table 2 for stereotaxic coordinates) and fixed to the skull with a liquid bonding resin (Superbond, Sun Medical) and dental acrylic cement (Unifast).

### Odor-reward associative learning

Go/No-go operant conditioning task was performed using custom-built olfactometers as described previously (Alonso *et al*., 2012; Mazo *et al*., 2022). At least 3 weeks after the last stereotaxic injection, mice were water-deprived (maintained at ∼85% of their initial body weight) and progressively trained to receive odors in a sampling port and then get a water reward from a waterspout 5 cm left from the odor port. Mice weight was closely monitored, and task rules training was performed without odor presentation. Once mice were able to stay at least 1.2 seconds in the odor port (presence detected by an infrared laser beam), odor association was allowed to proceed. For all trials, mice needed to enter the odor port and stay for at least 1 second before receiving an odor.

Mice were presented with a pair of odors in a pseudo-random fashion (no more than 3 times the same odor in a row), and only one (S+) was followed by the obtention of a water-drop, triggered by licking the spout, while the other one (S-) did not receive reinforcement of any kind. Odor was presented in the port for a maximum of 2 seconds, or stopped when mice removed their snout from the sampling port, which then left them 2 seconds to get the reward. If mice licked the spout after receiving S+, the result is a Hit, and a False Alarm (FA) after S-. If they did not lick after S+, result is a Miss, and a Correct Rejection (CR) after S-. The inter-trial interval was 5 seconds. Mice were tested on discrimination for 10 blocks of 20 trials, containing each 10 rewarded (S+) and 10 unrewarded odors (S-), with a 4 to 6 μL water drop as a reward.

A pseudorandom protocol was used during behavioral experiments to assign animals to the different olfactometers (six olfactometers in total). A given animal was never trained more than two consecutive days in the same device.

For each 20-trial block, performance is calculated as a percentage according to the following formula:

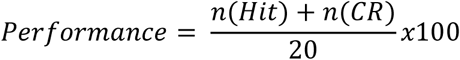

A threshold performance of 85% was required to consider that mice had correctly learnt to assign the reward value to the S+ and the non-reward value to the S−. The number of blocks needed to reach this criterion level was counted as the number of blocks employed before reaching a block with 85% correct responses.

For reversal, mice underwent 2 blocks of the initial rule, which were followed by 5 blocks where initial S+ became unrewarded while S- was rewarded. If a given mouse did not reach the 85% criterium in the first two blocks, the results of the reversal task were discarded.

Twenty-four hours or 2 weeks after last reinforced learning, memory was assessed by two blocks of 20 trials where previously learnt odors were presented but not rewarded. Memory score was then calculated according to:

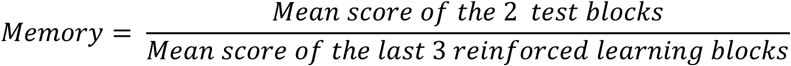

Automated olfactometer allow precise measurement of the time of events during the task. The odor-sampling time, called Reaction Time (RT), is measured between the arrival of the odor and the moment the mouse removes its head from the odor port, while First Lick (FL) is calculated between the end of RT and the first detected lick on the waterspout.

Odors were presented using glass vials containing the following couples: Hexanol/Octanal (first initial training), Amylacetate/Ethylbutyrate, (+)-Limonene/ (-)-Limonene. All odors were pure monomolecular odorants (Sigma-Aldrich), diluted in odor-less mineral oil (Sigma-Aldrich) at 1/100. Odors were generated by passing a 120mL /min stream of air over the surface of diluted odorants in disposable 50mL glass tubes. The odorant vapor was mixed with 3.20 L/min clean air before its introduction into the sampling port. Thus, the odor concentration delivered was 4.3% of the head space above the liquid odorant.

### Optogenetic experiments

To provide photo-stimulation for optogenetics excitation experiments, mice were tethered to the flexible optic fibers patch cords (Doric Lenses) coupled to a DPSS laser (473 nm, 150 mW, CNI Lasers) via a custom-built fiber launcher and controlled by the PS-H-LED laser driver connected to a custom-made pulse generator triggered by the automated olfactometer. Laser onset was synchronized with the closing of the diversion valve, allowing bilateral stimulation during odor presentation (both S+ and S-) with 473nm (output fiber intensity: 5mW) laser at 33Hz (5ms pulse duration) for 500 msec. The absolute light intensity was calibrated before each experiment. Animals in which fibers were outside of the brain target were discarded for the analysis (Mice n = 1/17).

### Electrophysiology

Mice were deeply anesthetized with intraperitoneal injection of Ketamine (100mg/Kg) and Xylazine (10mg/Kg) and swiftly decapitated. The OB and frontal cortices were rapidly dissected and placed in ice-cold ACSF containing in mM: 124 NaCl, 3 KCl, 2.8 MgSO4, 26 NaHCO3, 1.25 NaHPO4, 20 glucose, 0,5 CaCl2 (∼310 mOsm, pH 7.3 when bubbled with a mixture of 95% O2 and 5% CO2; all chemicals from Sigma France). They were then glued to a block of 4% agarose and placed, submerged in ice-cold ACSF, in the cutting chamber of a vibrating microtome (Leica VT 1200S). Horizontal slices (300 μm thick) of the OBs were placed in bubbled ACSF in a warming bath at 35°C for 30 min and then at room temperature (*i.e.*, 22 ± 1°C).

For whole-cell recordings, individual slices were placed in a chamber mounted on a Zeiss Axioskop upright microscope, and continuously perfused (1.5 mL/min) with 30°C ACSF containing in mM: 124 NaCl, 3 KCl, 1.3 MgSO4, 26 NaHCO3, 1.25 NaHPO4, 20 glucose, 2 CaCl2 (∼310 mOsm, pH 7.3 when bubbled with a mixture of 95% O2 and 5% CO2; all chemicals from Sigma France) (Warner Instrument inline heater). Slices were visualized using a 40x water-immersion objective. We obtained whole-cell patch-clamp recordings from visually targeted TdTomato-labeled GCs. Patch pipettes, pulled from borosilicate glass (OD 1.5mm, ID 0,86mm, Sutter Instrument, UK; P-97 Flaming/Brown micropipette puller, Sutter Instrument Co, UK), had resistances of 6–10 MΩ and were filled with a Cesium-Methane sulfonate-based solution (in mM: 126 Cs-MeSO3, 6 CsCl, 2 NaCl, 10 HEPES, 0.2 Cs-EGTA, 0.3 GTP, 2 Mg-ATP, 280–290 mOsm, pH 7.3). All membrane potentials indicated in the text are corrected for a measured liquid junction potential of +10mV. Labelled cells were identified by the presence of TdTomato in the tip of the patch pipette after membrane rupture. Recordings were obtained via an Axon Multiclamp 700B. Signals were filtered at 2 kHz and sampled at intervals of 20–450 μs (2.2–50 kHz) according to the individual protocols. Series resistance (Rs), and membrane resistance (Rm) were estimated using peak and steady-state currents, respectively, observed in response to a 5mV membrane step. Currents mediated by Na^+^ voltage-gated channels were measured under voltage-clamp conditions. Depolarizing pulses (100 ms) from –70 mV to incremental steps (5 mV), up to +10 mV, were given at a rate of 1 Hz. Na+ currents were measured after subtraction of scaled passive current responses to the appropriate voltage steps. IPSCs were recorded at Vc = 0 mV and EPSCs at Vc = -70 mV.

Synaptic events were elicited by photo-activation of ChR2^+^ axon terminals stimulation using a 470 nm light-emitting diode (LED; Xcite by Lumen Dynamics) illuminating the sample trough the objective. Duration of the light pulses was adjusted (from 0.1 to 6,4ms) for each cell to evoke minimal to maximal PSCs. Data were acquired using Elphy software (Gerard Sadoc CNRS; Gif-sur-Yvette, France) and analyzed with Elphy and IgorPro (Neuromatic by Jason Rothman).

### Calcium imaging using fiber photometry

During behavioral tasks, neurons infected with GCamp8f expressing AAV vector were continuously excited with a 473-nm solid-state laser (10mW; Crystal Lasers) via a 430-μm multimode optical fiber (output intensity < 0.1 mW). The emitted fluorescence was collected by the same fiber, filtered through a dichroic mirror and a GFP-emission filter (452–490 nm/505–800 nm; MDF-GFP, Thorlabs), filtered (525 ± 19 nm) and then focused on a NewFocus 2151 Femtowatt photodetector (Newport). Blue light reflected in the light path was also filtered and measured with a second amplifying photodetector (PDA36A; Thorlabs). The signals from the two photodetectors were digitized by a digital-to-analogue converter (Micro1401-3 A/D interface, CED) at 5000 Hz and then recorded using Spike2 software (CED, UK). Paxinos Brain atlas was used as reference to check viral vector injection and fiber optic implantation site in each animal. Mice where injections or optic fiber were outside were discarded for the analysis (n = 3/7).

### Immunolabelling

Mice were deeply anesthetized with intraperitoneal injection of ketamine (150mg/Kg) and Xylazine (10mg/Kg) and were intracardially perfused first with Saline (0.9% NaCl), then with 4% paraformaldehyde in 0.1M phosphate buffer. Brains were removed, post-fixed 2 hrs. in 4% PFA (except for VGAT staining) and cryoprotected in PBS-Azide 0.02 %-sucrose 30% overnight. Sagittal slices (60μm thick) were obtained with a microtome and were stored with PBS-Azide 0.02%. Immunostaining was performed on free-floating sections. Non-specific staining was blocked in PBS-Triton (PBST) 0.5% with 10% Normal Goat Serum for 1h30 at room temperature followed by incubation in primary antibody solution (see Table 3) in PBST 0.5% over 1 or 2 nights at 4 °C. Sections were subsequently washed 3 times in PBST 0,5% (for 15 min each) and then transferred to secondary antibody solution containing a DNA-specific fluorescent probe (Hoechst; 1:5000) in PBST 0.5% for 2 hours at room temperature. Sections were again washed 3 times in PBST (for 15 min each) before being mounted onto glass slides and coverslipped using Fluoromount-G mounting media.

**Table 3.** Antibodies.

| Antibody | Provider | Dilution | Figure |
| --- | --- | --- | --- |
| Chicken anti-GFP | Abcam (ab13970) | 1/1000 | S1, S2, 5, S6 |
| Mouse anti-RFP | Rockland (200-301-379) | 1/1000 | 5, S6 |
| Rabbit anti-vGAT | Synaptic System (131002) | 1/1000 | 5, S6 |
| Goat anti-Chicken 488 | Invitrogen (A11039) | 1/1000 | S1, S2, 5, S6 |
| Goat anti-Mouse 568 | Invitrogen (A21134) | 1/1000 | 5, S6 |
| Goat anti-Rabbit 647 | Invitrogen (A21244) | 1/1000 | 5, S6 |

For VGAT puncta analysis, images of slices were obtained using a confocal laser-scanning microscope (LSM 980, 63x oil objective; Zeiss®) in stack of optical slices (0.36 μm thick along the Z axis). Images acquisition and quantification were performed blindly to the brain slice condition.

### VGAT puncta image analysis

Image analyses were performed using the software Imaris®. Surfaces were created for both eYFP (GABAergic fibres) and TdTomato (adult-born GCs) using a minimum surface size of 2500 voxels. VGAT-immunoreactive puncta were detected using the *Spots* function with an estimated diameter of 1 µm. VGAT spots were then filtered based on proximity to both the eYFP and tdTomato surfaces; spots were retained if they were within 0.5 µm of the eYFP surface and within 0.5 µm of the tdTomato surface. Retained VGAT spots were classified into four compartments based on their location: soma, proximal dendrite (<30 µm from the soma), distal dendrite (>30 µm from the soma within the granule cell layer, GCL), or apical dendrite (located in the external plexiform layer, EPL). To quantify total GABAergic fiber signal, the mean gray value of the eYFP channel was measured in each z-stack and averaged across all stacks for a given image. VGAT puncta density was quantified as the total number of VGAT spots divided by the number of z-stacks.

### Fiber photometry analysis

Analysis of fluorescence signals was performed using custom-written MATLAB codes (Mathworks, MATLAB R2023b). Raw GCaMP8f fluorescence signals were first pre-processed: traces were down sampled from 5000 to 50 Hz and smoothed over a window of 0.02 s. Bleaching effects were controlled by fitting a two-term exponential, and signals were adjusted accordingly only if the R-square of the goodness of fit was above 0.6. After that, we computed a z-score defined as [F(t) – mean(F(t))]/sd(F(t)) with the mean and standard deviation calculated on a sliding window of 100 s. The different trial outcomes were then identified and aligned to the photometry signals. For graphical representation in the peri-stimulus time histograms, the signal from each trial was normalized based on the 2 s preceding the opening of the final valve. Then, using the same normalization, we calculated the Area Under the Curve (AUC) divided by its duration for several time intervals: (1) the 1 s interval during which the odor is prepared, (2) the variable interval during which the mouse has its nose in the odor port, (3) the 2 s interval after the mouse removed its nose from the port, (4) the 2 s interval after the preceding one. The latency and the value of the GCaMP peak from the odor onset were also calculated for each trial.

### Statistical Analyses

Statistical analyses were performed with GraphPad Prism 9.0 (except for calcium imaging analysis). Most of the tests used in this study are non-parametric tests (Mann-Whitney test, Kruskal-Wallis test with Uncorrected Dunn’s test for multiple comparisons, Friedman test) due to the low sampling and non-normal distribution of data. When a parametric test was performed (Unpaired t-test, One-way ANOVA), normality was verified using the Kolmogorov-Smirnov test. Differences were considered significant for p < 0.05. Outliers were identified with ROUT column analyses and were removed for Q ≥ 0.5%. Calcium imaging analysis for each odor pair were performed using R 4.5. Mixed models were used taking into account mouse ID, training day and trial number as random effects, and followed by Tukey pairwise comparison.

### Data and code availability

The datasets and codes that support the findings of this study are available from the corresponding author upon reasonable request.

Further detailed method and material information can be found in the Supplemental information.

## INTRODUCTION

Adult neurogenesis, the production of new neurons in the mature brain, occurs throughout the entire life in most mammalian species (Ming & Song, 2011; Lepousez *et al*., 2015). In rodents, this process primarily occurs in the hippocampus and olfactory bulb (OB) and confers structural and functional plasticity to neural networks, thereby contributing to learning and memory (Kempermann, 2022), emotional regulation (Alonso *et al*., 2024), and the modulation of cognitive decline associated with aging (Alonso-Moreno *et al*., 2025).

In the olfactory system, neuroblasts are generated in the subventricular zone, migrate to the OB through the Rostral Migratory Stream (RMS) and differentiate mostly into granule cells (GCs) once fully mature (Carleton *et al*., 2003). Furthermore, young abGCs demonstrate enhanc(Lledo & Valley, 2016). Consistent evidence has shown that adult-born neurons possess unique morphological or electrophysiological properties compared to those generated during the neonatal period (Dejou *et al*., 2024). Functionally, adult-born granule cells (abGCs) acquire the ability to generate action potentials later than neonatally-born granule cells (nnGCs), but they exhibit heightened excitability ed synaptic plasticity (Nissant *et al*., 2009) and unique adaptive capabilities in response to sensory manipulations (Magavi *et al*., 2005; Saghatelyan *et al*., 2005; Mandairon *et al*., 2018; Forest *et al*., 2019). They also display distinctive features in their synaptic output (Valley *et al*., 2013). Importantly, although some of these unique characteristics disappear when their maturation is fulfilled (Bardy *et al*., 2010), several studies have shown that mature abGCs keep long-term specific features not found in their neonatal counterparts (Mizrahi, 2007; Valley *et al*., 2013).

In addition to their unique cellular properties, the specific contribution of abGCs might also stem from their distinctive connectivity. At the OB network level, abGCs are predominantly located in the deeper regions of the granule cell layer (GCL) compared to nnGCs (Lemasson, 2005), which may impact their network effects. Selectively depleting abGCs significantly reduces the inhibition of mitral cells (MCs) by dendrodendritic connections and impairs gamma oscillation generation in the OB (Breton-Provencher *et al*., 2009).

Several studies have already linked abGCs to olfactory perception, including odorant detection (Breton-Provencher *et al*., 2016) and discrimination (Enwere *et al*., 2004), olfactory innate behaviors (Sakamoto *et al*., 2011; Garrett *et al*., 2015; Muthusamy *et al*., 2017), perceptual learning (Moreno *et al*., 2009), olfactory learning (Alonso *et al*., 2012; Grelat *et al*., 2018), odor fear memory (Valley *et al*., 2009), but also short-term (Rochefort *et al*., 2002; Breton-Provencher *et al*., 2009) and long-term olfactory memory (Lazarini *et al*., 2009; Sultan *et al*., 2010; Alonso *et al*., 2012).

Perception does not rely solely on feedforward transfer of information from sensory organs; it is also shaped by feedback from higher-order brain areas that provide contextual information and predictions. Accordingly, top-down signals modulate early sensory processing, with corticofugal projections conveying context and predictive information to lower-level regions. These fibers are involved in selective attention, object expectation, and associative features, highlighting their involvement in learning and memory processes (Rao & Ballard, 1999; Gilbert & Li, 2013; Keller & Mrsic-Flogel, 2018).

In the olfactory system, the anterior piriform cortex (APC) and the anterior olfactory nucleus (AON), which together form the anterior olfactory cortex (AOC), send dense innervation back to the OB. Glutamatergic projections target all main types of OB neurons and induce di-synaptic inhibition onto MCs and Tufted Cells (TCs) (Gao & Strowbridge, 2009; Boyd *et al*., 2012; Markopoulos *et al*., 2012). These reciprocal connections between the OB network and the cortex are important for proper OB oscillations (Martin *et al*., 2006; Kay, 2014) and decorrelation of OB output activity (Otazu *et al*., 2015). They also modulate odor perception threshold (Soria-Gómez *et al*., 2014), odor-association learning (Wu *et al*., 2020; Chae *et al*., 2022; Chen *et al*., 2022; Hernandez *et al*., 2025) and social transmission of odor preference (Wang *et al*., 2020). GCs, and particularly abGCs, are at the converging endpoint of both widespread top-down inputs to the OB and bottom-up olfactory inputs mediated by M/TCs (Lepousez *et al*., 2013). In line with this idea, we demonstrated that abGCs appear to play a pivotal role in the coding of odor value during olfactory learning. Exposure to reward-associated odors increases the activity of abGCs but not preexisting ones. Notably, activating abGCs when a rewarded odor is presented enhances discrimination learning and improves the ability to update odor value during a reversal association (Grelat *et al*., 2018).

Indeed, olfactory learning strengthens corticofugal fibers synapsing onto immature abGCs, increasing both excitatory and inhibitory inputs in the deep dendritic domain that preferentially receives cortical projections. Selective optogenetic stimulation of glutamatergic cortical projections to the OB confirmed that learning potentiates these inputs onto abGCs (Lepousez *et al*., 2014). Cortical excitatory inputs are essential for abGCs plasticity during learning: blocking these inputs prevents learning-induced plasticity, while their artificial stimulation mimics plasticity without odor exposure. This plasticity is specific to the abGCs activated by odor exposure during learning (Wu *et al*., 2020). Thus, diverse top-down control and learning-mediated connectivity changes may underlie the functional differences between adult- and neonatal-born GCs.

For decades, it was thought that cortico-bulbar projections were exclusively glutamatergic. However, we recently uncovered a direct inhibitory projection from the olfactory cortex to the OB. These fibers primarily target interneurons in the GCL (GCs and dSACs) and output neurons (MCs and TCs). They provide inhibition of the spontaneous and odor-evoked activities of these neurons. Moreover, stimulating cortical GABAergic fibers increases beta oscillation power in the OB, while silencing these fibers using pharmacogenetics impairs the discrimination of similar odor mixtures (Mazo *et al*., 2022). However, how these fibers selectively impact abGCs is still unknown.

In the present study, we investigated whether long-range cortical GABAergic connections are recruited in the OB during olfactory learning and whether these fibers exhibit specific connectivity with mature abGCs. Using *in vivo* calcium imaging via fiber photometry, we showed that GABAergic fibers are activated with both odor- and reward-dependent dynamic upon presentation of odor cues. Optogenetic activation of these fibers impaired both the learning and the reversal of an odor-reward association. Importantly, by using *ex vivo* patch-clamp recordings coupled with an olfactometer-driven olfactory learning task, we demonstrated that olfactory learning potentiates cortical GABAergic inputs onto abGCs, but not their neonatal counterpart nor glutamatergic inputs.

## RESULTS

### GABAergic top-down projections from the anterior olfactory cortex are activated during an odor-reward associative learning

To investigate the potential contribution of centrifugal fibers originating from the anterior olfactory cortex (AOC), we measured the activity of these fibers in behaving animals. To this end, we injected a Cre-dependent GCaMP8f-expressing viral vector into vGAT::Cre mice, targeting the posterior part of the anterior olfactory nucleus (AONp) in order to restrict the virally-transduced area, and implanted an optic fiber above the olfactory peduncle (**Figure 1A**). This approach allowed us to perform fiber photometry calcium activity recordings specifically of passing cortico-bulbar inhibitory projections while mice performed an olfactory discrimination learning task.

**Figure 1.**
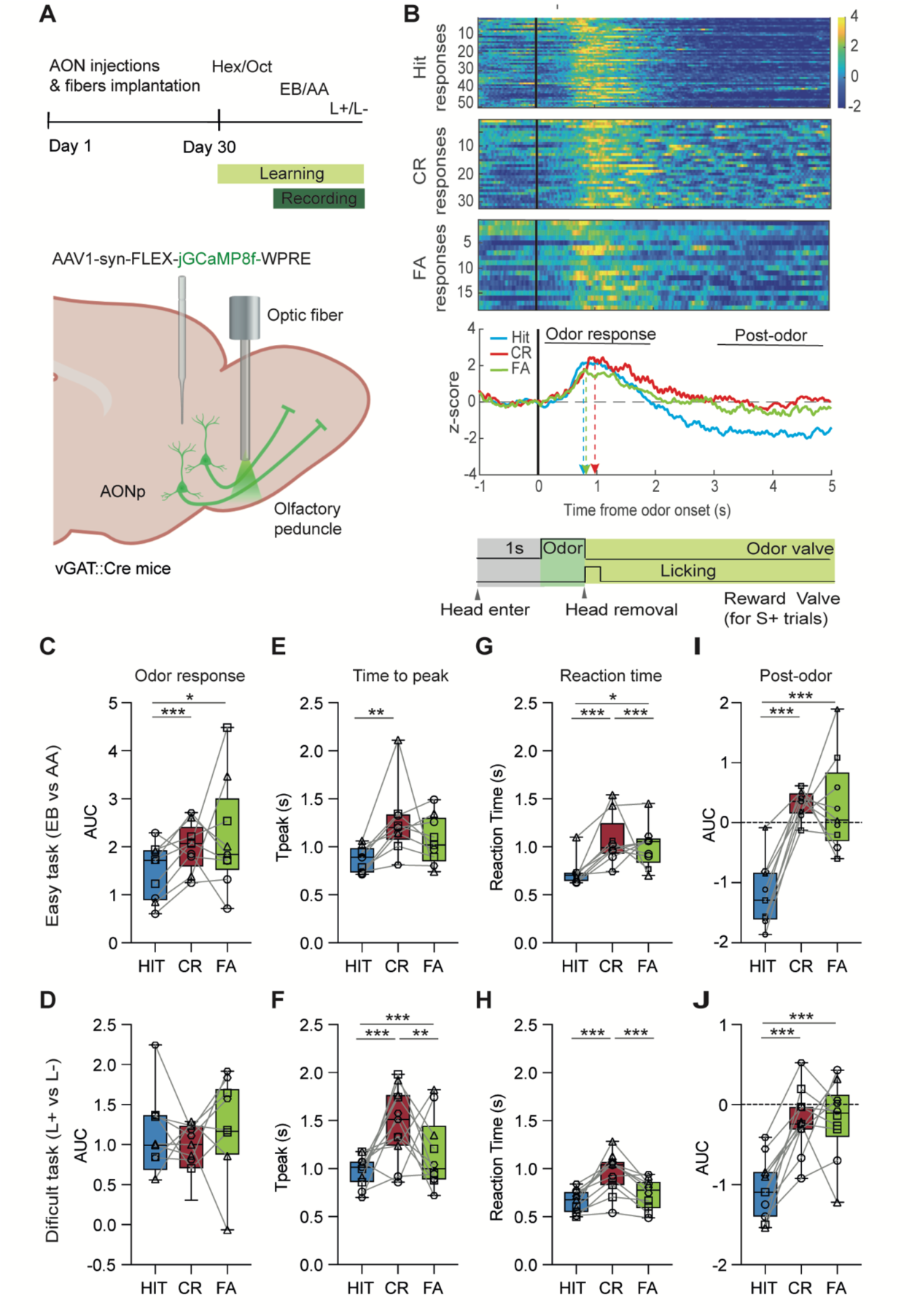
AONp-to-OB GABAergic projections are activated during Go/No-go task. **A**. Upper panel: Timeline of the experiments. Lower panel: Stereotaxic injection of GCaMP8f vector in AONp of VGAT::Cr mice and optic fiber implantation above the olfactory peduncle to allow recording of fibers projecting to the OB. **B**. Upper panel: Heatmaps representing all individual trials z-score for “Go’’ correct response after S^+^ (HIT responses), “no-go” correct responses to S^-^ (CR responses) and ‘’Go’’ incorrect response after S^-^ (FA responses). Middle panel: Averaged traces for HIT (blue), CR, (red) and FA (green). Lower panel: Diagram of the trial timeline. **C-D** Summary graph of the area under the curve (AUC) during odor responses (0-2s) for EB vs AA (**C;** (Mixed-effect model Χ^2^ = 48.37969, p<0.0001) and L+ vs L- (**D**; Mixed-effect model Χ^2^= 3.348253, p = 0.18747). **E-F**. Summary graph of the time between the onset of odor delivery and the maximal Z-score value (time to peak) for EB vs AA (**E;** (Mixed-effect model Χ^2^ = 99.31759, p<0.0001) and L+ vs L- (**F**; Mixed-effect model Χ^2^ = 93.25118, p<0.0001). **G-H**. Summary graph of the time between the onset of odor delivery and the head removal from the odor port (reaction time) for EB vs AA (**G;** (Mixed-effect model Χ^2^ = 216.1429, p<0.0001) and L+ vs L- (**H**; Mixed-effect model Χ^2^ = 210.2016, p<0.0001). **I-J**. Summary graph of the area under the curve (AUC) during post odor response (2-4s) for EB vs AA (**I;** (Mixed-effect model Χ^2^ = 428.9851, p<0.0001) and L+ vs L- (**J**; Mixed-effect model Χ^2^ = 192.4263, p<0.0001). Data are represented as box plot from mean value per animal per day of training, for HITs (blue), CRs (red) and FA (green) for each odor pair (EB: Ethylbutyrate; L+: limonene +; L-: limonene -). *p<0.05, **p<0.01, ***p<0.001.

For that, we used an operant conditioning Go/No-go task in custom-built olfactometers, in which the mice learned to discriminate successively three reinforced odors associated with a water reward (positive stimulus, S+) against non-rewarded odors (negative stimulus, S−; see **Figure S1**). Animals were trained to discriminate respective odor pairs until reaching the performance criterion of 85% for 3 different pairs of odors (**Figure S1**).

We observed that odor presentation activates AONp GABAergic projections and that this activity persists after animals withdraw from the odor port (**Figure 1B**). In an easy discrimination task (Ethyl-butyrate vs. Amyl acetate, EB vs AA), the mean area under the curve (AUC) of the odor-evoked response was higher for S⁻ odor responses (CR: correct rejections; FA: false alarms) than for S⁺ odor response (HIT) (**Figure 1C**). Notably, MISS responses to S+ were extremely rare in our protocol. A similar activation pattern was observed in the difficult task (Limonene⁺ vs. Limonene⁻, L^+^ vs L^-^); however, AUC did not differ between S⁺ and S⁻ odors-evoked responses, likely due to the high similarity of the enantiomers used in this task (**Figure 1D**).

The peak of maximum activity during CR responses was significantly delayed compared to HIT and FA responses in both tasks (**Figure 1E, F)**. At the behavioral level, mice exhibited longer reaction times (intervals between odor arrival and port withdrawal) when exposed to non-rewarded odors compared to rewarded ones (**Figure 1G,H**), consistent with our previous findings (Grelat *et al*., 2018). To assess the relationship between time-to-peak and reaction time, while accounting for mouse-specific effects and response type, we employed a mixed-effects statistical model. Although the relationship was significant, the effect size was modest: 7.71% for EB vs. AA (95% CI: 2.97–12.89) and 3.47% for L⁺ vs. L⁻ (95% CI: 0.51–7.80). These results suggest that reaction time differences (and thus odor exposure duration may only partially explain the observed kinetic disparities in odor responses.

Following water reward delivery in HIT responses (2–4 s post-odor onset), activity was significantly reduced compared to CR and FA responses, a pattern consistent across all analyzed tasks (**Figure 1I, J**). Collectively, our findings demonstrate that GABAergic top-down projections from the AONp to the OB are odor-activated during associative learning and exhibit distinct temporal dynamics depending on the behavioral outcome triggered by the olfactory stimulus.

### Optogenetic disruption of AONp GABAergic neurons impairs olfactory learning and reversal, but not memory

We next sought to manipulate GABAergic corticobulbar fibers to probe whether changes in the activity of these inputs are sufficient to alter behavior. To address this, we used optogenetics to specifically activate the GABAergic fibers of the AONp during odor presentation, through the expression of ChR-YFP protein (ChR2 group) in vGAT::CRE mice. Control mice were only injected with GFP-expressing AAV (control group; **Figure 2A, B**). For specific *in vivo* manipulation of neurons projecting to the OB, all the mice were implanted bilaterally with optic fibers on top of the olfactory peduncles, and 4 weeks later trained in olfactory learning tasks as previously described (**Figure 2C)**. In the first initiation task without light stimulation, both control and ChR2 groups performed similarly, which allowed us to discard potential off-target effects of viral vector expression (**Figure 2D, Laser OFF**). Once animals reached the criterion, they were trained in the same task while receiving a light stimulation beginning at the odor onset at 33Hz and lasting 500 msec for both S+ and S-stimuli (**Figure 2C, D, Laser ON**). This duration was set considering the mean reaction time measured for this task equivalent to odor exposure that triggers GABAergic corticobulbar fiber activation (see **Figure 1G, H).** No effect was observed once a discrimination rule was already acquired, demonstrating that stimulation protocol does not disturb discrimination acuity or behavioral performance.

**Figure 2.**
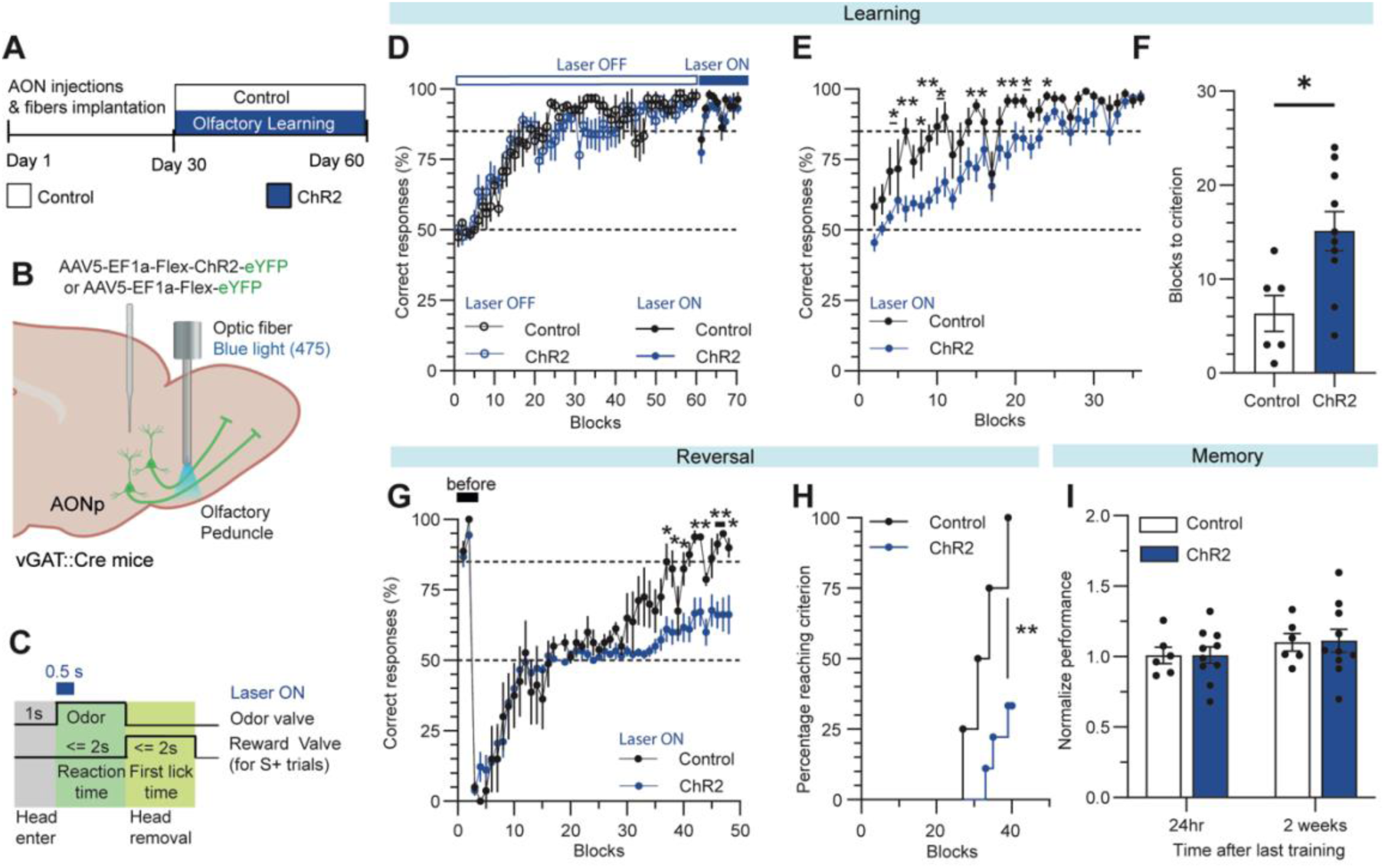
Optogenetic manipulation of AONp-to-OB GABAergic projections disrupts odor-reward association. **A**. Timeline of the experiments. **B**. Stereotaxic injection of ChR2 vector in AONp of VGAT::Cre mice and optic fiber implantation above the olfactory peduncle to allow stimulation of fibers projecting to the OB.**C**. Timeline of light stimulation in each trial. **D.** Learning performance of a discrimination task (Hexanol vs Octanal, 1/100) without light stimulation (Laser OFF, white band; Mixed-effect analysis, group: F_(1,14)_ = 0.4539, p=0.5115, block F_(6.623, 85.12)_ = 18.89, p<0.0001; interaction: F_(6.623, 85.65)_ = 1.184, p= 0.3212) and during trials when light was on (Laser ON, blue band, 33Hz, 5ms pulse duration; Mixed-effect analysis, group: F_(1,14)_ = 2.679, p=0.1239, block F _(4.433, 61.56)_ = 5.797, p=0.0003; interaction: F_(4.433, 61.56)_ = 0.6286, p= 0.6598, n=6-10). **E**. Learning performance in discrimination task ((+)-Limonene vs (-)-Limonene) (Mixed-effect analysis, group: F_(1,14)_ = 7.574, p=0.0107, block F_(6.620, 92.49)_ = 23.02, p<0.0001; interaction: F_(6.620,92.49)_ = 1.771, p= 0.1065, n= 6-10). **F**. Number of blocks necessary to reach the 85% performance criterion in limonene learning task (t-test, p=0.0130). **G**. Discrimination performance during reversal of the limonene learning task (S- becomes rewarded and S+ is no longer rewarded). (Mixed-effect analysis, group: F_(1,11)_ = 4.864, p=0.0496, block F_(4.744,51.98)_ = 43.26, p<0.0001; interaction: F_(4.744,51.98)_ = 3.005, p= 0.0203, n= 4-9). **H.** Cumulative curve of mice reaching the 85% criterion in reversal task across time (Mantel-Cox test, p=0.0068, n=4-9). **I**. Mean performance for 40 test trials (No reward, no stimulation), 24hr (left) and 2 weeks (right) after the last training session (2-way ANOVA, group: F_(1,14)_ = 0.0047, p=0.9494, time F_(1,14)_ = 3.871, p = 0.0693; interaction: F_(1,14)_ = 0.0075, p= 0.9320 n= 6-10). Data are shown as mean ± SEM and individual data points. *p<0.05, **p<0.01, ***p<0.001.

Moreover, stimulation of AONp GABAergic feedback impair the olfactory learning discrimination of a new odor pair (**Figure 2E**), reflected by a significant increase in the number of blocks necessary to reach the criterion (**Figure 2F**). Interestingly, no difference was apparent in reaction time or first lick between groups (**Fig S2**). This result suggests a predominant role of GABAergic inputs in the early phases of the odor–reward association. To challenge this idea, animals were trained in a reversal version of the task. Mice had to reverse the values associated with an odor pair that they had already learnt. For this, the odorant previously associated with a reward was unreinforced and previous unrewarded odor was now associated with a reward (**Figure 2G**). As expected, during the first few blocks of the reversal task, all animals performed below the chance level, as they kept following the previous odor–reward association rule. Both groups similarly reset the odor-reward association by performing again at 50% chance level performance after few blocks (block to reach 50% after reversal in Control: 9,25 ± 2.56 and ChR2: 7,77 ± 0.89 Mann-Whitney test p = 0.7021, n = 4-9; **Figure 2G**). However, the adjustment to the new odor–reward association was strongly affected by GABAergic fibers activation in ChR2 group compared to control (**Figure 2G**). Most animals in ChR2 group never managed to reach the success criterion with the new rules in at least 40 blocks, as shown when percentage of animal reaching criterion were compared between groups (**Figure 2H**).

We also analyzed whether light activation altered olfactory memory. For these experiments, animals were trained until reaching success criterion while GABAergic fibers were activated as previously described. Then, mice were tested for odor memory recall 24h then 2 weeks after the end of the training session without light stimulation. The data was normalized to last blocks performance during training to rule out differences due to learning acquisition. Memory was not altered in the ChR2 group with respect to control mice, showing that these fibers do not seem necessary for olfactory memory encoding (**Figure 2I**). Collectively, these results demonstrate that the activation of GABAergic corticobulbar fibers projecting to the OB participates in odor-reward association and cognitive flexibility.

### Adult-born granule cells receive learning-sensitive GABAergic inputs from the anterior olfactory cortex

We have demonstrated that abGCs strongly contribute to odor-reward association during discrimination learning (Alonso *et al*., 2012; Grelat *et al*., 2018) and our previous work showed that GABAergic corticobulbar projections innervate both output and local OB neurons (Mazo *et al*., 2022). However, it remains unknown whether long-range cortical GABAergic projections preferentially target abGCs and whether this cortical feedback undergoes learning-dependent changes.

To study the functional connectivity of corticofugal inputs on abGCs vs nnGCs, we labelled GCs progenitors generated at specific ages of mice using stereotaxic injections of TdTomato-expressing lentiviral vectors in the RMS of pups at postnatal day 6 (P6) or adult 2-month-old mice (P60) (**Figure 3A**). One month later, we injected an adeno-associated virus (AAV) expressing channelrhodopsin-2 (ChR2) fused to green fluorescent protein (YFP) into the anterior olfactory cortex (AOC; see **Figure 3A**) to target both glutamatergic and GABAergic cortico-bulbar projections. Importantly, the ChR2-YFP cells were exclusively located in the AOC, and we did not detect direct viral diffusion from the injection site to the bulb. Additionally, injection into the AOC did not label M/TCs retrogradely nor significantly labelled migrating neuroblasts *en route* to the OB (28 YFP+ GCs/mm3 equivalent of 1,1x10^-4^ % of total neurons in GCL based in (Parrish-Aungst *et al*., 2007); n=14), and a negligible number of cells in the HDB/MCPO regions were observed (82 GFP+ cells/6633 ChAT^+^ analyzed, from n=18 mice). This study concentrates only on mature newly-born interneurons by starting experiments at least 8 weeks post injection (wpi), when these neurons are already integrated in the bulbar circuit and their density in the GCL have reached a plateau (Bardy *et al*., 2010).

**Figure 3.**
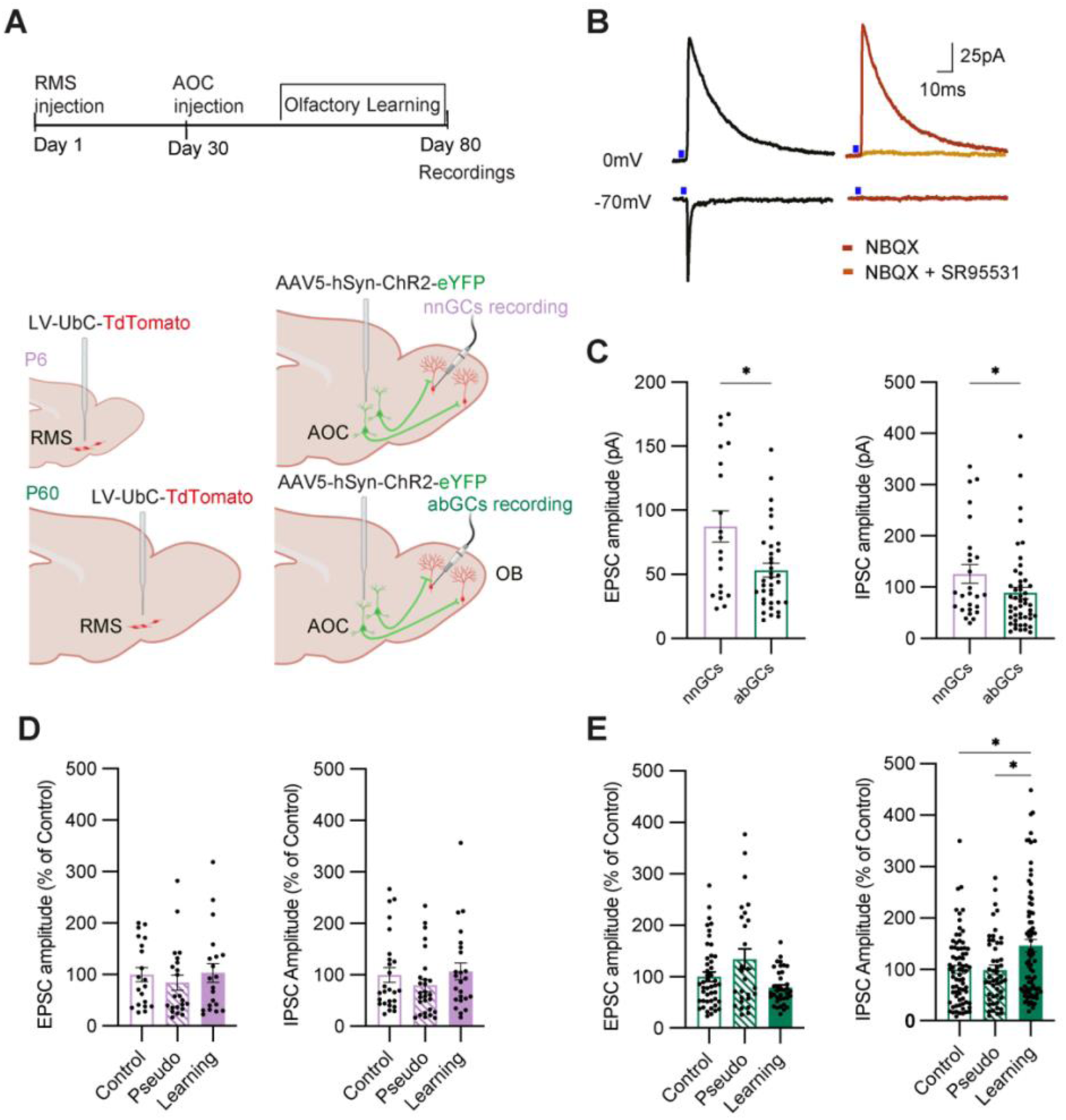
Associative learning tunes cortical inputs on adult-born granule cells. **A.** Upper panel: Chronology of stereotaxic injections and electrophysiology recordings. Lower panel: Stereotaxic injections of TdT-expressing lentiviral vector in the Rostral Migratory Stream (RMS) at neonatal (P6) or adult (P60) age and of ChR2-expressing virus in the anterior olfactory cortex (AOC). **B.** Representative mean traces of light evoked post-synaptic currents at various voltages in absence of ionotropic receptors blockers (black traces) or in presence of 10µM NBQX (red trace) or 10µM NBQX + 10µM SR95531 (yellow traces). **C**. Left: Light-evoked EPSCs amplitude (Mann-Whitney test, p=0.0446, n=20-46). Right: Light-evoked IPSCs amplitude (Man-Whitney test, p=0.0134, n=26-39). **D**. Left: Amplitude of light-evoked EPSCs on neonatally born GCs (nnGCs, Kruskal-Wallis test, p=0.5387, n=20-22-20). Right: Amplitude of light-evoked IPSCs on nnGCs (Kruskal-Wallis test, p=0.2993, n=26-27-24). **E.** Left: Amplitude of light-evoked EPSCs on adult-born GCs (abGCs, Kruskal-Wallis test, p=0.2235, n=46-27-37). Right: Amplitude of light-evoked IPSCs on abGCs (Kruskal-Wallis test, p=0.0196; Dunn’s test, Control vs Pseudo, p=0.9615; Control vs Learning, p=0.0137; Pseudo vs Learning, p=0.0237, n=73-50-80). For **D** and **E**, Electrophysiology data is expressed as % of control mean value. Data is shown as mean ± SEM and individual data points, *p<0.05.

We used whole-cell patch-clamp recordings of visually targeted Td-tomato^+^ GCs and activated ChR2^+^ cortical terminals using full field illumination with 470nm light through the microscope objective. We recorded light-evoked excitatory and inhibitory postsynaptic currents (PSCs) in the voltage-clamp mode (**Figure 3B**). We did not find any difference in membrane resistance when comparing cells generated at neonatal and adult age (nnGCs: 831MΩ ± 88.3 n- = 21; abGCs: 923.7MΩ ± 83.4 n = 29; Mean ±SEM, Mann-Whitney test, p = 0.572), but we observed a higher amplitude of voltage-gated Na^+^ current in adult-born neurons, in agreement with previous results (Carleton *et al*., 2003) (nnGCs: 1266 ± 108.3 n = 29; abGCs: 1827pA ± 172 n = 26; Mean ±SEM, Mann-Whitney test, p = 0.0109). We isolated AMPAR- and GABA_A_R-mediated currents by changing the membrane potential and using pharmacology to isolate specific and monosynaptic responses (**Figure 3B**). For inhibitory post-synaptic currents (IPSCs), we analyzed only those that were resistant to the application of 10 µM NBQX to ensure that they originated directly from AOC projections rather than resulting from a polysynaptic circuit (Mazo *et al*., 2022). As expected, SR95531 (a GABAA receptor antagonist) abolished all inhibitory synaptic responses.

The amplitude of light-evoked EPSCs and IPSCs was greater in nnGCs than in abGCs (**Figure 3C**). Together, these findings indicate that nnGCs form stronger connections with—or receive more inputs from—the AOC compared to abGCs. This suggests that top-down fibers differentially innervate the two populations of GCs, potentially giving rise to distinct microcircuit properties and functions.

Next, we investigated the impact of olfactory learning on both P60 and P6-injected groups. All mice learned to discriminate successively three reinforced odors associated with a water reward (Learning group; **Figure S3**). For these experiments, the go/no-go task was also performed in one group with a randomly given reward, which prevented the association with a specific odorant but exposed mice to the same odorants and reward (called pseudo-learning group). Animals were exposed daily to a pair of odorants (200 trial per day for 21 days). Mice from the learning group showed a rapid learning of discrimination, gradually reaching the performance criterion of 85% for 3 different pairs of odors (**Figure S3**), while pseudo-learning ones remained at the 50% chance level, as expected. For both neonatal and adult-born groups, performances were similar in terms of the learning rate (**Figure S3**). Meanwhile, control mice were not exposed to odors, nor to reward, and were kept in their home cage throughout the experiment. Once all learning tasks were completed and 24 h after the last training session, patch-clamp recordings were performed. When recording nnGCs, no changes were observed in light-evoked excitatory and inhibitory responses in both learning and pseudo-learning groups compared to control mice (**Figure 3D**). For abGCs recording, no difference was found in the amplitude of EPSCs (**Figure 3E, left**). However, we showed that amplitude of IPSCs significantly increased on abGCs cells after olfactory learning in respect to both pseudo-learning and control animals (**Figure 3E, right**). Overall, our data show that an olfactory associative learning increases inhibitory corticofugal inputs on abGCs, but not on their neonatal counterparts.

### Adult-born granule cells display more inhibitory synapses with cortical terminals after learning

We then decided to focus on the increase of inhibition observed after odor-reward association in adult-born neurons. To specifically label and manipulate GABAergic feedback, we employed again a conditional labelling approach. We used the same injection protocol as previously described, but this time using a conditional AAV-Cre-dependent-ChR2-YFP in vGAT::Cre mice to specifically label GABAergic AOC cells (**Figure 4A**) and we performed the same learning paradigm (**Figure S4**).

**Figure 4.**
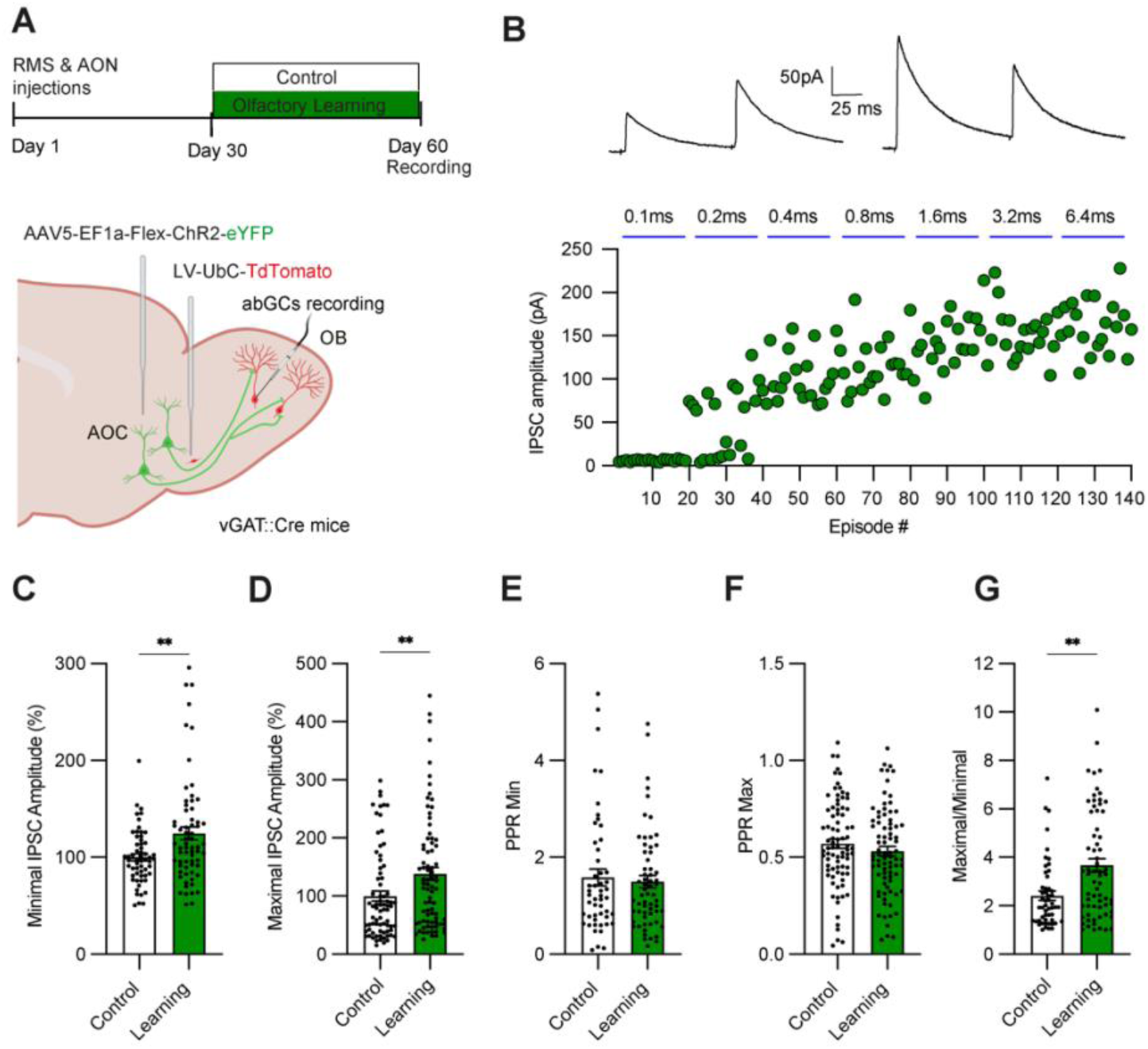
Associative learning enhances cortical GABAergic inputs onto adult-born granule cells through postsynaptic potentiation. **A**. Upper panel: Chronology of stereotaxic injections and electrophysiology recordings. Lower panel: Stereotaxic injections of TdT-expressing lentiviral vector in the Rostral Migratory Stream (RMS) at adult age and of ChR2-expressing virus in the anterior olfactory cortex (AOC) in VGAT::CRE mice. **B**. Upper panel: Representative average traces of minimal and maximal light-evoked IPSCs (Paired-pulse stimulation; ISI= 100ms). Lower panel: Representative example of progressive stimulation intensity and associated IPSCs amplitudes **C.** Minimal light-evoked IPSCs amplitude (Mann-Whitney test, p=0.0017, n=61-67). **D.** Maximal light-evoked IPSCs amplitude (Mann-Whitney test, p=0.0202, n=75-77). **E**. Paired-pulse ratio for minimal IPSCs (Mann-Whitney test, p=0.898, n=51-62). **F**. Paired-pulse ratio for maximal IPSCs (Mann-Whitney test, p=0.232, n=82-81) **G.** Maximal/minimal ratio (Mann-Whitney test, p=0.0019, n=52-66). Data is shown as mean ± SEM and individual data points, **p<0.01.

To refine our results, we isolated minimal and maximal responses in the recorded GCs by using incremental light pulse duration (from 0.1 to 6.4ms) (**Figure 4B**). Minimal responses were collected at the shortest time duration eliciting synaptic currents. Maximal responses were collected when the amplitudes of the responses reached a plateau. We found that both minimal and maximal light-evoked responses were increased by olfactory learning (**Figure 4C, D**), confirming and extending our previous data on WT mice. Interestingly, this potentiation was not observed when recording light evoked IPSCs onto MCs (**Figure S5**). In order to assess the locus of the potentiation, we performed paired-pulse stimulation (ISI 100ms) and measured paired-pulse ratio (PPR) as a proxy of the presynaptic probability of GABA release. We found no difference in PPR between groups (**Figure 4E, F**), which is in favour of a post-synaptic mechanism of potentiation. Finally, the Max/Min ratio was also higher in these cells (**Figure 4G**), suggesting that learning could promotes the formation of a greater number of AOC inhibitory synapses onto abGCs.

To probe the morphological correlates of these electrophysiological changes, we performed histology on fixed slices to count the number of synapses according to their location on the abGCs cellular domains. We used the same injection protocol in the RMS with 1/10 viral vector dilution to allow the detection of individual neuron structure (**Figure 5A**). We observed that GABAergic cortical top-down axonal boutons (in green), with most labeling in the GCL and lesser expression in the internal plexiform layer (IPL) and mitral cell layer (MCL), made putative synapses on TdTomato+ abGCs (in red; **Figure 5B**). In addition, to validate the presumptive inhibitory inputs on abGCs, we labelled and quantified the presence of vesicular GABA transporter (VGAT, in grey, **Figure 5B**), a presynaptic protein essential for GABA accumulation into synaptic vesicles.

**Figure 5.**
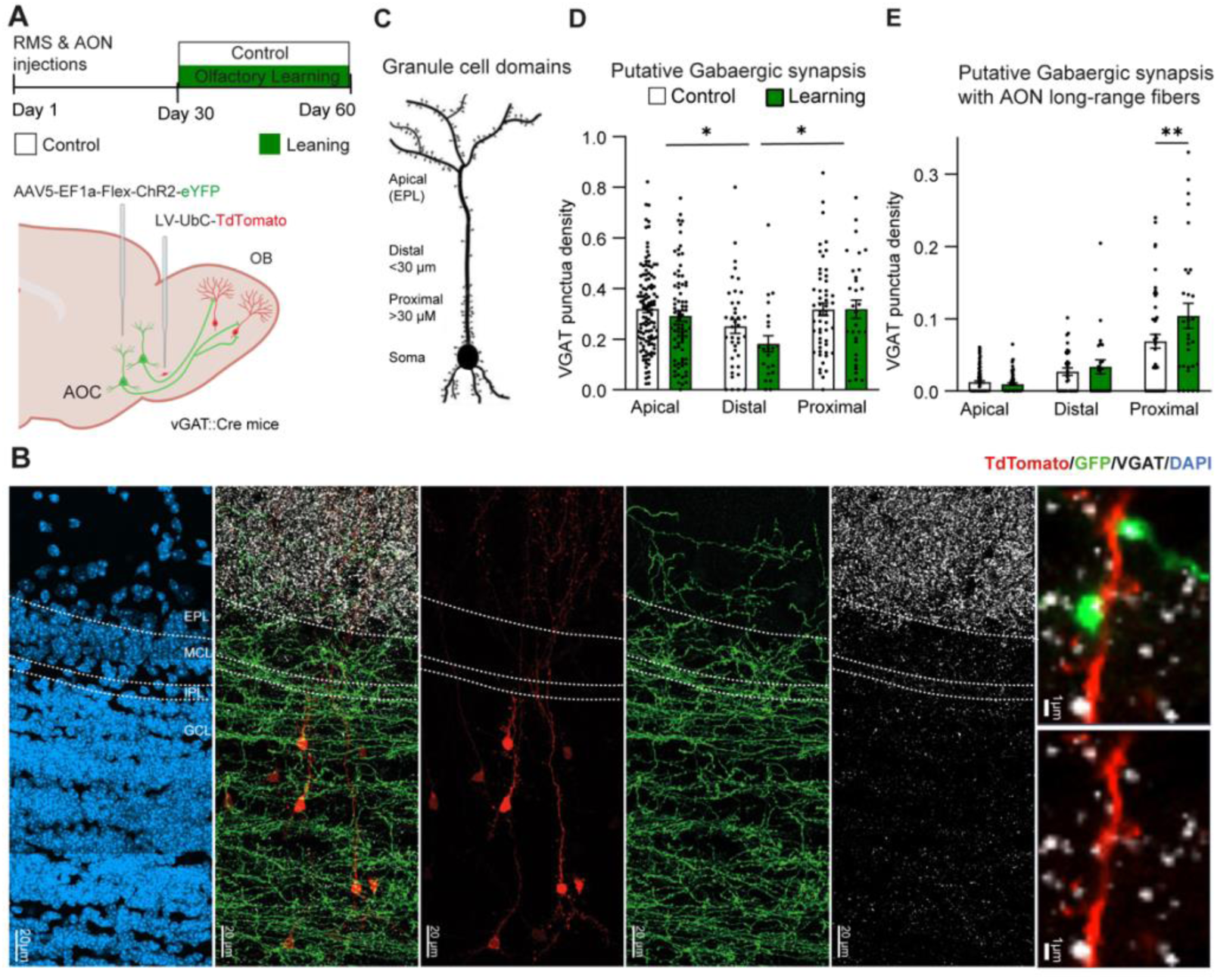
Adult born granule cells display more inhibitory synapses with AONp-to-OB GABAergic fiber after learning. **A**. Upper panel: Timeline of the experiments. Lower panel: stereotaxic injections of TdT-expressing virus in the Rostral Migratory Stream (RMS) at adult age and ChR2-expressing virus in the Anterior Olfactory Cortex (AOC) in VGAT::CRE mice. **B**. Left: confocal image showing staining at different OB layers. Right: Zoomed in view of confocal image obtained after immunolabeling of adult-born GCs (Tdtomato, red), cortical inhibitory afferences (eYFP, green) and VGAT synaptic protein (grey). **C**. Schematic representation of GCs anatomical segmentation. **D**. Density of VGAT+ puncta (grey) in the different dendritic domains of abGC (red) (n=23-117 dendritic segments per condition, n=3-6 mice); 2-way ANOVA, group: F_(1,337)_ = 2.131, p=0.1453; domain F_(2,337)_ = 6.785, p=0.0013; interaction: F_(2,337)_ = 0.6400 p=0.5280; Apical vs Distal = p=0.0024; Apical vs Proximal = 0.9396; Distal vs Proximal = 0.0028. **E**. VGAT+ puncta (grey) colocalized with GABAergic projecting axons (green) in different dendritic domains of abGC (red); (n=23-111 dendritic segments per condition, n=3-6 mice); 2-way ANOVA, group: F_(1,313)_ = 4.685, p=0.0312; domain: F_(2,313)_ = 75.83, p<0.0001; interaction: F_(2,313)_ = 4.970, 0.0075; Apical p=0.09249; Distal p=0.9323; Proximal p=0.0020). Density is expressed as puncta per µm. Data are shown as mean ± SEM and individual data points, *p<0.05, **p<0.01.

First, we analyzed the distribution of presumptive inhibitory inputs along the dendritic tree and soma of GCs. For that, GC dendritic trees were subdivided into four compartments: somatic and proximal, distal, and apical dendritic domains (**Figure 5C**). The proximal compartment was defined as the first 30μm of the main dendrite starting from the soma, the distal compartment as the adjacent segment starting 30μm away but still within the GCL, and the apical compartment as the dendritic arbor located in the EPL. In our samples, basal dendrites were often difficult to visualize – probably due to a reduced expression of TdTomato in this compartment – and were not included in our analysis.

Our results showed that putative GABAergic inputs (vGAT+/TdTomato+) are located throughout the dendritic compartments of abGCs, although they are significantly less abundant on distal in respect to apical and proximal domains. Moreover, no differences were found between groups suggesting that learning does not change the total amount of GABAergic synapses impinging on mature abGCs (**Figure 5D**). However, when we analyzed the number of VGAT+ puncta colocalizing with corticobulbar GABAergic fibers (vGAT+/TdTomato+/eYFP+), we found a significant global increase in learning group in respect to control group, mostly due to significant difference observed in proximal domain (**Figure 5E**). Concerning the somatic compartment, no difference was found in the number of total putative GABAergic synapsis (control soma: 0,02087 ± 0,001926; Learning soma: 0,02136 ± 0,002501, puncta per µm^2^, p=0,9244, Mann-Whitney test, n = 42-64 soma per condition, 4-6 mice), as well those colocalizing with corticobulbar GABAergic fibers (control soma: 0,005420 ± 0,0009974, n = 64; Learning soma: 0,004565 ± 0,001068, puncta per µm^2^, p = 0,6453; Mann-Whitney test, n = 42-64 soma per condition, 4-6 mice). Importantly, several histological controls rule out that the difference found could be due to sampling or immunostaining bias (**Figure S6**).

Our data validated the observed functional changes and suggested that odor-reward association increases the number of GABAergic synapses arriving from cortical long-range inputs onto abGCs.

## Discussion

In this study, we found that cortical GABAergic fibers exhibit behavior-dependent activation in response to odor cues by using *in vivo* calcium imaging. Optogenetic activation of this GABAergic circuit impaired both the acquisition and reversal of an odor-reward association, highlighting its critical role in olfactory learning. Ultimately, we demonstrated that these cortical projections form synaptic connections with abGCs that are sensitive to olfactory learning.

Analyzing the dynamics of GABAergic fiber activation *in vivo* using Ca²⁺ fiber photometry during an olfactory task could provide valuable insights into their functional role. Our recordings revealed a rapid increase in fiber activity, beginning few milliseconds after odor onset and persisting throughout odor presentation. This response peaked on average one second after odor onset—a time point at which the animal had typically already withdrawn from the odor port. Given the kinetics of GCaMP8f, we speculate that the signal begins to decay around the time of port departure. While this observation is purely correlative, the most parsimonious explanation is that the response decline coincides with odor offset. However, we cannot exclude the possibility that this activity contributes to the animal’s decision to terminate odor sampling. Furthermore, we observed a pronounced suppression of fiber activity during HIT responses, that coincided with water reward delivery. This suppression was unrelated to the animal’s performance or the timing within the training session. Furthermore, we didn’t observe a similar pattern during FA response in which the animal also decides to lick but receives no reward. This decrease in calcium activity could reflect a reduction of the cortical inhibitory tone upon reward presentation. Alternatively, water intake *per se* may induce some changes in animal’s breathing pattern during licking behavior which could participate in the decrease of cortical inhibitory tone.

To assess more directly their functional role, we manipulated the activity of GABAergic fibers using optogenetic stimulation during an olfactory task. In our experiments, we delivered 33 Hz stimulation for 500 ms at odor onset, across all trials (both S+ and S– presentations). This manipulation impaired learning and disrupted task reversal, a finding that aligns with previous work demonstrating that direct optogenetic activation of abGCs produces the opposite effect—enhancing learning and improving reversal performance in a similar task (Grelat *et al*., 2018). An alternative interpretation is that our stimulation imposed an artificial, non-physiological activation pattern, interfering with the natural dynamics of fiber recruitment that normally occur during odor presentation. Specifically, by homogenizing responses to S+ and S– stimuli, we may have disrupted the differential excitatory/inhibitory balance of the cortical drive between rewarded and unrewarded odors, and thus alter odor discrimination learning. Notably, once the learning criterion was reached without photo-stimulation, we observed that neither abGC (Grelat *et al*., 2018) nor GABAergic activation did not further modify discrimination acuity of an already established odor–reward association. These results suggest a predominant role of abGCs and GABAergic top-down inputs in the early phases of the odor–reward association. Importantly, our approach lacked cellular specificity, as the stimulation broadly targeted both abGCs and nnGCs, as well as those that may or may not respond to the presented odor. Additionally, off-target effects of cortical axon stimulation on other postsynaptic cell types, such as MCs or short-axon cells (SACs), cannot be ruled out.

The present report demonstrates that abGCs exhibit heightened inhibitory responses to centrifugal fiber stimulation following associative olfactory learning but no change in excitatory responses. Previous studies have reported increased cortical excitation onto abGCs as a result of olfactory learning (Lepousez *et al*., 2014; Wu *et al*., 2020). The discrepancy between these findings and our present results can be attributed to several methodological differences. As we mentioned before, the cells analyzed in both previous studies were younger (32 to 35 days post-injection) compared to those in our experiments (60-80 days post-injection). Additionally, Wu (Wu *et al*., 2020) and colleagues utilized *in vivo* two-photon microscopy to record Ca2^+^ activity in GCaMP-positive axon terminals while in our study we analyzed post-synaptic currents evoked by photoactivation of ChR2^+^ axon terminals using whole-cell patch-clamp recordings in acute OB slices. Interestingly, in our previous study we already showed that the density of putative GABAergic synapses (Gephyrin^+^ puncta) was increased after olfactory learning, specifically on the proximal domain of GCs dendrites, as well as the frequency of spontaneous IPSCs (Lepousez *et al*., 2014). At the time, the existence of GABAergic projections from the cortex had not yet been described, and thus had not been specifically studied. It is possible that a similar increase in cortical GABAergic fibers after learning occurs both in young (30-35 dpi; Lepousez *et al*., 2014) and more mature abGCs (60-80 dpi; present study), which is not the case for cortical glutamatergic synapses that change only in young cells (Wu *et al*., 2020). Interestingly, no effect of learning was observed on inhibitory synapse arriving from the cortex to M/TCs, reinforcing the idea of a specific plastic changes on abGCs.

Our results suggest that the enhanced GABAergic response following learning may arise through at least two distinct mechanisms. First, we observed an increase in minimal IPSC amplitude without change in the paired-pulse ratio, a finding consistent with postsynaptic potentiation, such as an upregulation of GABA receptor density at the synapse. Second, the elevated Max/Min amplitude ratio, combined with histological evidence, indicates an increase in the number of synapses between GABAergic top-down projections and abGCs after learning. However, since the temporal dynamics of centrifugal GABAergic synaptogenesis during abGCs maturation remain unclear, it is challenging to determine whether learning promotes *de novo* synapse formation or prevents the pruning of existing synapses (Whitman & Greer, 2007).

Upon comparing the synaptic properties of nnGCs with those of abGCs, we observed a greater amplitude of both glutamatergic and GABAergic currents evoked by cortical inputs on nnGCs. Interestingly, these findings suggest that weaker centrifugal inputs in abGCs may provide a broader dynamic range, thereby facilitating their potentiation during learning processes. This finding contrasts with a previous study that reported stronger glutamatergic connections in abGCs (Wu *et al*., 2020). As we already mentioned, the recorded abGCs in that study were only 2 weeks old, and the control group could have consisted of cells generated at various time points, resulting in a highly heterogeneous population. In contrast, our study compares mature abGCs (60-80 days post-injection), that are already fully integrated in the circuits (Bardy *et al*., 2010), with nnGCs generated specifically at postnatal day 6 (P6). Additionally, the enhanced excitation observed in abGCs relative to nnGCs was evaluated by measuring the total charge resulting from high-frequency light stimulation over a 1-second period, whereas our analysis assessed individual post-synaptic currents to short light pulses. These methodological differences preclude a direct comparison of our findings on centrifugal excitation with those reported in the aforementioned study.

What would be the functional consequences of enhancing inhibitory responses specifically in abGCs? Our findings, together with previous studies, indicate that both excitatory and inhibitory cortical inputs are predominantly integrated within the proximal dendritic domain (<100 µm)—a strategic location for regulating the initiation and propagation of action potentials (APs) through the dendritic arbor (Pressler & Strowbridge, 2019). Notably, excitatory cortical inputs onto GCs have been shown to gate dendro-dendritic inhibition of MCs (Balu *et al*., 2007; Restrepo *et al*., 2009). Based on this, we hypothesize that an increase in GABAergic input could moderate this drive, thereby further refining the tuning of MCs output. However, prior research has demonstrated that learning strengthens abGCs responses to learned odors (Breton-Provencher *et al*., 2009; Grelat *et al*., 2018; Mandairon *et al*., 2018) and enhances abGC-to-MC connectivity (Huang *et al*., 2016). This suggests that learning induces odor-specific plasticity in abGCs, which may then provide targeted inhibition to responsive MCs. In this context, increased inhibition onto “non-responsive” abGCs could serve to reinforce plasticity mechanisms within odor-specific abGC populations, thereby sharpening the neural representation of learned odors.

Alternatively, we can hypothesize that centrifugal inhibition primarily functions to refine the temporal dynamics of abGC activity, thereby modulating network processing. GABAergic long-range projections are thought to contribute to the generation of oscillatory rhythms in the brain, facilitating the synchronization of distant neural regions. For instance, stimulation of basal forebrain GABAergic neurons has been shown to enhance theta and gamma oscillations upon local activation of their terminals in the glomerular layer (GL) and granule cell layer (GCL), respectively (Villar *et al*., 2021). Furthermore, 33 Hz stimulation of olfactory cortex inhibitory fibers increased beta-range oscillations within the OB network (Mazo *et al*., 2022), a phenomenon previously linked to learning-related processes involving centrifugal fibers (Martin *et al*., 2004).

This study focuses on long-range GABAergic inputs, primarily originating from the AON (Mazo *et al*., 2022). As a key early olfactory processing area, the AON exhibits extensive reciprocal connections and has increasingly been linked to critical functions in both normal and pathological conditions (Brunert *et al*., 2023). Recent research has identified distinct cortical glutamatergic feedback loops mediated by OB outputs and their preferred cortical targets. Depending on their origin—whether from the PC or the AON—these loops have been involved in different functional roles (Chae *et al*., 2022).

In this line, another brain area that sends long range GABAergic projections toward the OB is the horizontal limb of the diagonal band of Broca (HDB), which selectively targets GCs, SACs, and specific subtypes of PGCs (Nunez-Parra *et al*., 2013; Case *et al*., 2017; Sanz Diez *et al*., 2019; Constantinescu *et al*., 2025). This pattern of innervation does not align with the one we described for cortical GABAergic fibers, supporting the possibility of an alternative function (Mazo *et al*., 2022). Furthermore, given the potential co-release of GABA and acetylcholine (ACh) from these projections, their functional impact on OB circuitry remains complex and context-dependent (De Saint Jan, 2022; Yu *et al*., 2025). Nevertheless, these fibers appear to contribute to the disinhibition of M/TCs, thereby facilitating the discrimination of similar odorants (Nunez-Parra *et al*., 2013). Recent evidence further suggests that HDB GABAergic projections exhibit dynamic activity patterns in response to reward-associated odors, increasing their activity during odor presentation and decreasing it below baseline upon reward delivery (Hanson *et al*., 2021). This dynamic closely parallels the activity patterns observed in cortical GABAergic fibers, as shown in the present study. More recently, a specialized subpopulation of calretinin-expressing (CR+) GABAergic neurons within the HDB was shown to selectively inhibit GCs and have a critical role in odor-associative learning, as chemogenetic inhibition of these neurons disrupts learning without affecting odor perception (Constantinescu *et al*., 2025).

Collectively, these findings emphasize the functional heterogeneity of long-range GABAergic pathways to the OB, where distinct circuits differentially modulate odor processing and learning. This highlights the nuanced roles of specific neuronal networks in olfactory cognition. In other sensory modalities, long-range GABAergic pathways flexibly tunes neocortical computation according to individual’s experience, preferentially targeting local interneurons (Schroeder *et al*., 2023). As authors mention, this disinhibitory connectivity provides a more flexible and dynamic substrate for circuit control than direct excitation supplied by classical top-down afferents. Further work is required to fully identify long-range GABAergic-GC loops and their specific role in olfaction.

In conclusion, this study provides important insights into two key areas: (1) the implication of adult neurogenesis in reward-associated learning and its distinct contributions to neural plasticity, and (2) the function of top-down inhibitory projections in integrating experience-dependent signals to modulate early sensory processing, particularly during odor discrimination. While further research is required to elucidate their precise mechanisms under varying internal states and pathological conditions, both neurogenesis and top-down projections are implicated in psychiatric disorders, including affective disorders (Alonso *et al*., 2024) and pathological aging (Culig *et al*., 2022). Given the significant impact of these disorders on sensory perception—particularly olfaction—understanding these pathways and their interplay may provide valuable insights into the pathophysiology of related symptoms.

## Acknowledgements

We thank Carine Moigneu for her assistance with some experiments, the Institut Pasteur Central animal facility for animal care and all Perception and Action lab members for their constructive inputs. This work was supported by Agence Nationale de la Recherche Grants ANR-15-NEUC-0004-02 “Circuit-OPL”, Laboratory for Excellence Programme “Revive” Grant ANR-10-LABX-73, ANR-AAPG2021 “EMOKET”, the Life Insurance Company AG2R-La Mondiale and Labex BioPsy. EP is funded by Ecole Normale Supérieure Paris-Saclay (CDSN); C-H De Badts is funded by Ecole Doctorale 158 (ED3C: Sorbonne Université, Université Paris Cité, Université Paris Sciences et Lettres).

## Disclosure

The authors declare no competing interests. The funding sources were not involved in study design, data collection, analysis and interpretation, writing or the decision to submit the article for publication.

## Author contributions

EP: Conceptualization, Methodology, Investigation, Formal analysis, Visualization, Writing – Original Draft, Writing – Review and Editing; CHDB, Fiber Photometry analysis, GL;SW: Conceptualization, Tool development; AH, JR, Investigation, Formal analysis; EJ: Statistical analysis of calcium imaging data, visualization; PML: Writing – Review and Editing, Supervision, Funding acquisition; MA, AN: Conceptualization, Methodology, Investigation, Formal analysis, Visualization, Writing – Original Draft, Writing – Review and Editing, Supervision, Funding acquisition.

---

**Supplementary figure 1.**
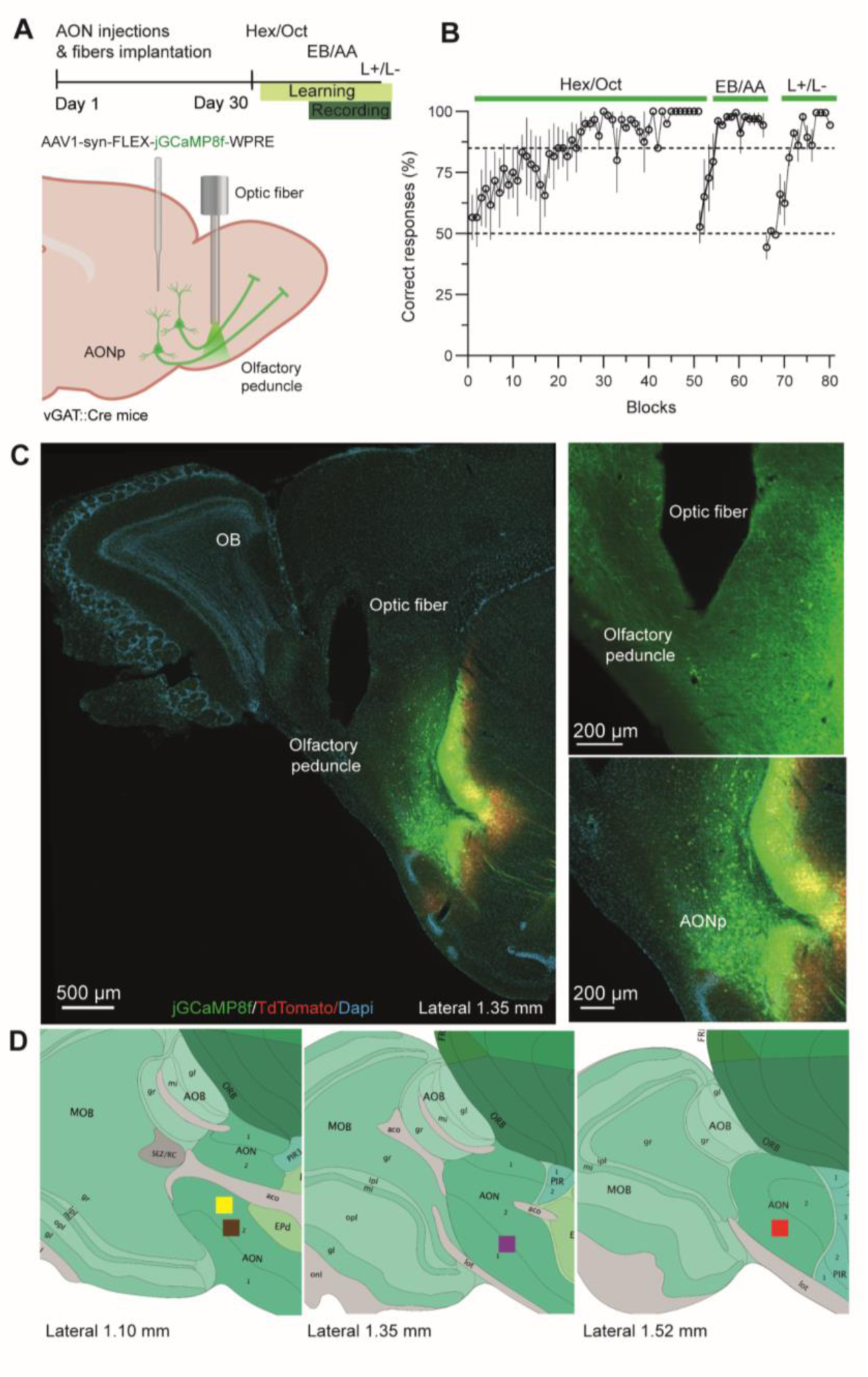
Associative learning performance throughout photometry experiments. **A**. Upper panel: Timeline of the experiments. Lower panel: Stereotaxic injection of GCaMP8f vector in AONp and optic fiber implantation above the olfactory peduncle to allow recording of fibers projecting to the olfactory bulb. **B**. Learning performance of three consecutive discrimination tasks (Hexanol vs Octanal; Ethyl Butyrate vs Amyl Acetate; (+)-Limonene vs (-)-Limonene). Data are shown as mean ± SEM. **C**. Representative image showing the location of fiber optic in the olfactory peduncle (left), detail of fiber optic tip showing recorded Gabaregic fibers (right top) and AONp injection site (right bottom). **D.** Implantation sites for recorded animals (hemisphere n=4; mice n=4).

**Supplementary figure 2.**
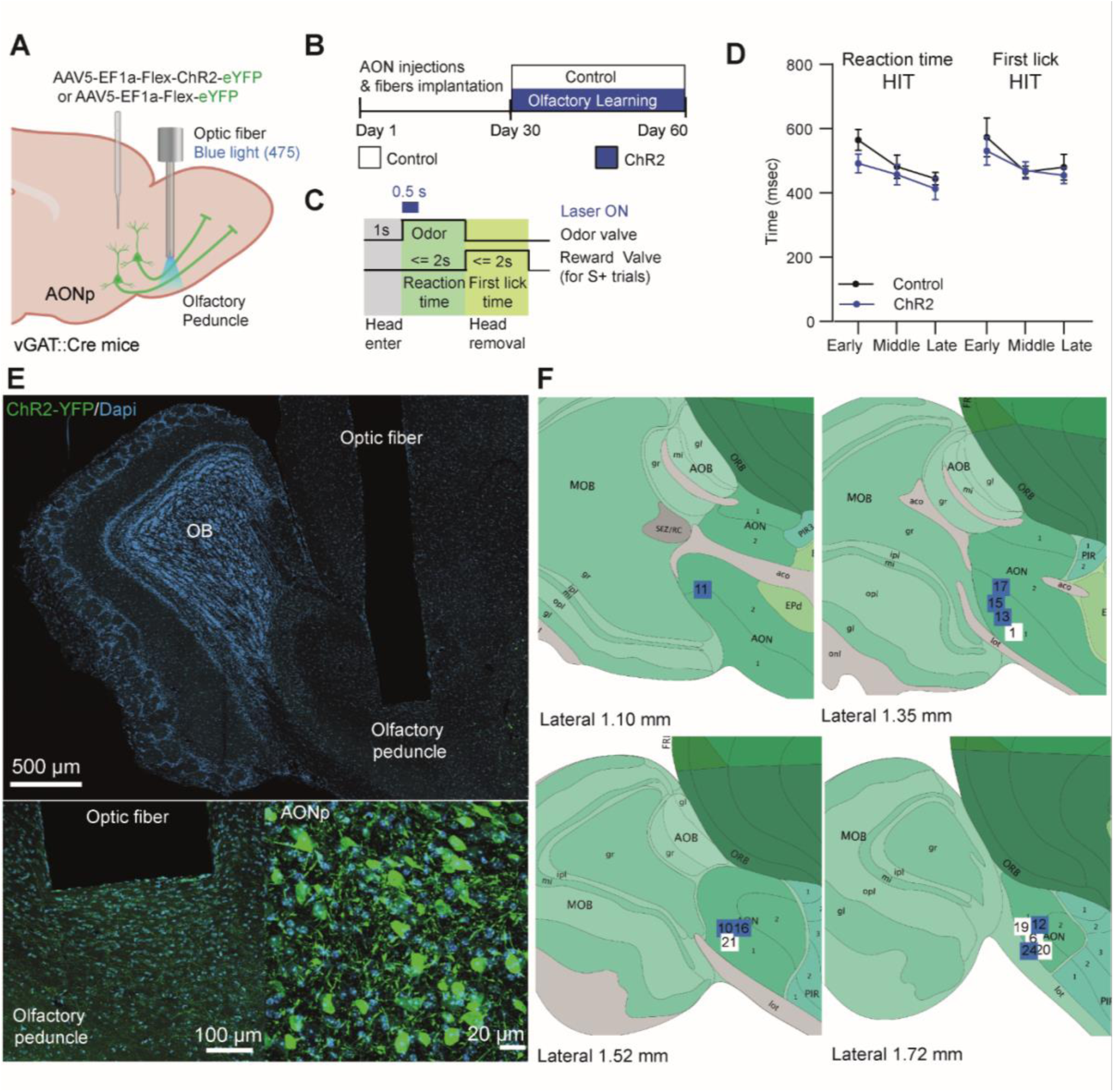
Activation of GABAergic fibers do not change reaction time. **A.** For optogenetic artificial activation of AONp-GABAergic projections, stereotaxic injection of ChR2 or control vector were performed in AONp and optic fibers were bilaterally implanted in the olfactory peduncle. **B**. Timeline of the experiments. **C**. Timeline of light stimulation in each trial. **D**. Activation of GABAergic projections do not change reaction time (Two-way RM ANOVA, group: F_(1,13)_ = 1.141, p=0.3049, learning phase F_(1.600, 20.80)_ = 13.42, p=0.0004; interaction: F_(2, 26)_ = 0.6286, p= 0.6598, n=6-9) nor movement time (Two-way RM ANOVA, group: F_(1,13)_ = 0.2202, p=0.6467, learning phase F_(1.420, 18.46)_ = 8.005, p=0.0061; interaction: F_(2, 26)_ = 0.6226, p= 0.4826, n=6-9) throughout learning . **E.** Representative image showing the location of fiber optics in the olfactory peduncle (top), detail of fiber optic tips showing light stimulated Gabaregic fibers (left bottom) and AONp injection site (right bottom). **F** Implantation sites for recorded animals (hemisphere n=10-18; mice n=5-10).

**Supplementary Figure 3.**
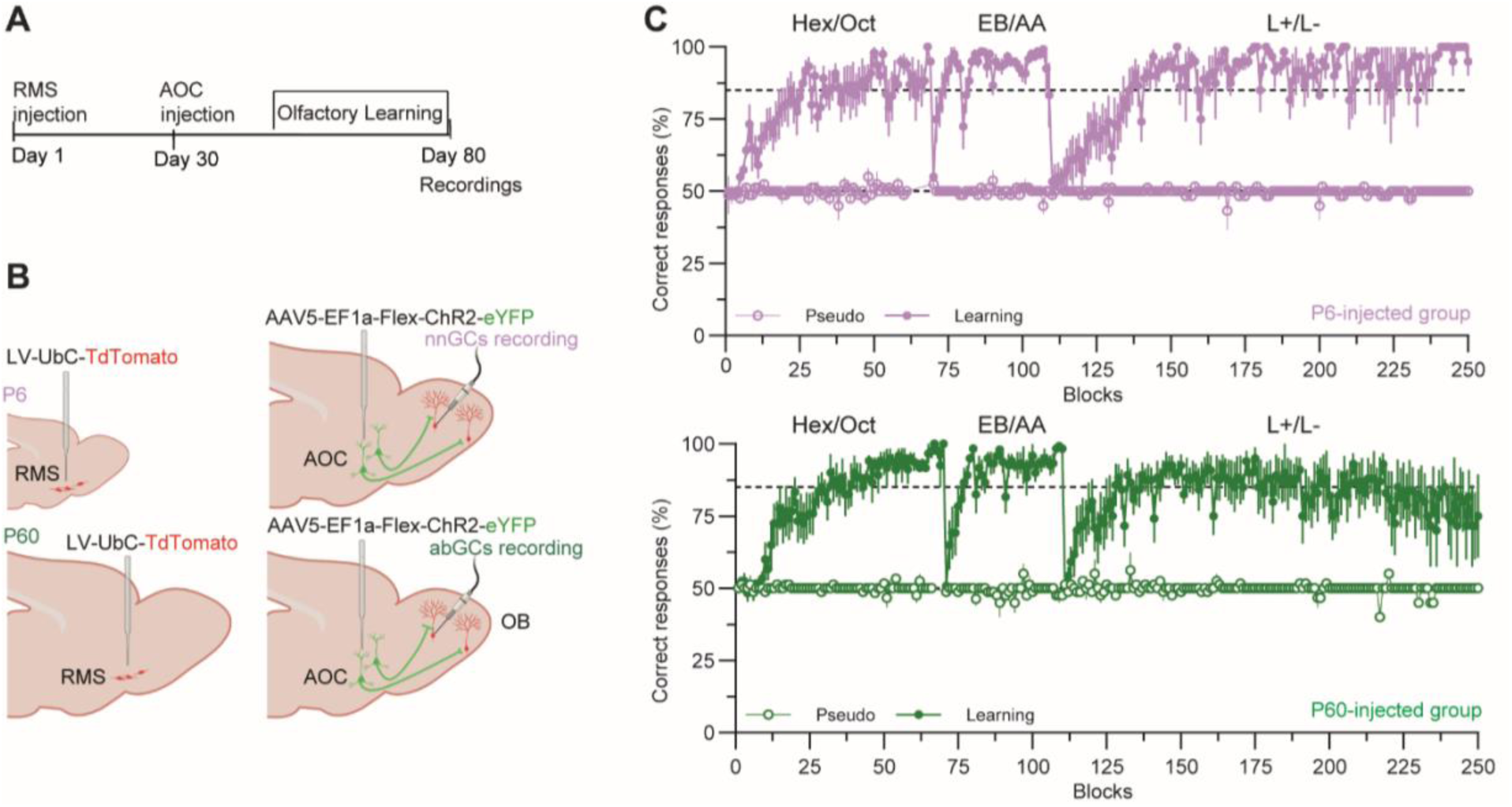
Associative learning performance of neonatal (P6) and adult (P60)-injected mice before patch clamp recordings. **A**. Chronology of stereotaxic injections and electrophysiology recordings. **B**. Stereotaxic injections of TdTomato-expressing lentiviral vector in the Rostral Migratory Stream (RMS) at neonatal (P6) or adult (P60) age and of ChR2-expressing virus in the anterior olfactory cortex (AOC). **C**. Learning performance of a three different discrimination tasks (Hexanol vs Octanal, 1/100; Ethyl Butyrate vs Amyl Acetate, 1/100; Limonene (+) vs Limonene (-), 1/100) of both learning and pseudo-learning group for P6-injected mice (pink, nbGCs labelled) and P60-injected mice (abCGs labelled, green). Data are shown as mean ± SEM.

**Supplementary Figure 4.**
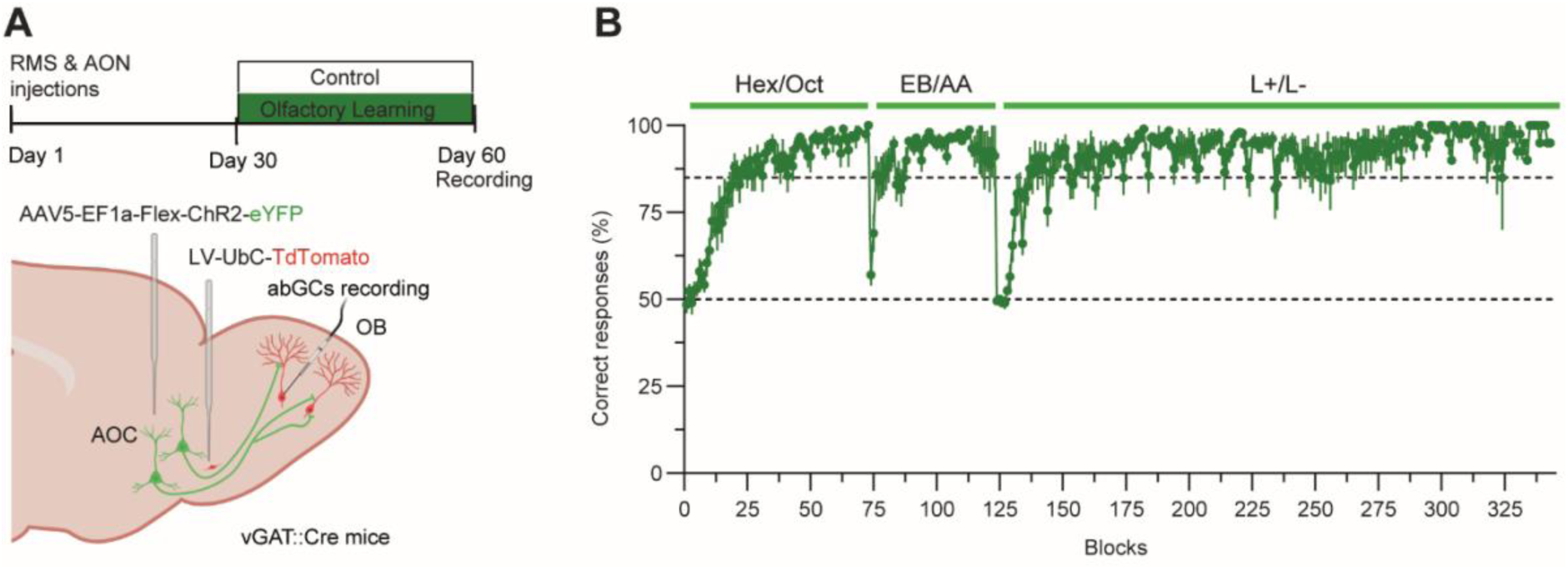
Associative learning performance of VGAT:CRE mice before patch clamp recordings. **A**. Upper panel: Chronology of stereotaxic injections and electrophysiology recordings. Lower panel: Stereotaxic injections of TdTomato-expressing lentiviral vector in the Rostral Migratory Stream (RMS) and of ChR2-expressing virus in the anterior olfactory cortex (AOC). **B**. Learning performance of a three different discrimination tasks (Hexanol vs Octanal, 1/100; Ethyl Butyrate vs Amyl Acetate, 1/100; Limonene (+) vs Limonene (-), 1/100). Data are shown as mean ± SEM.

**Supplementary figure 5.**
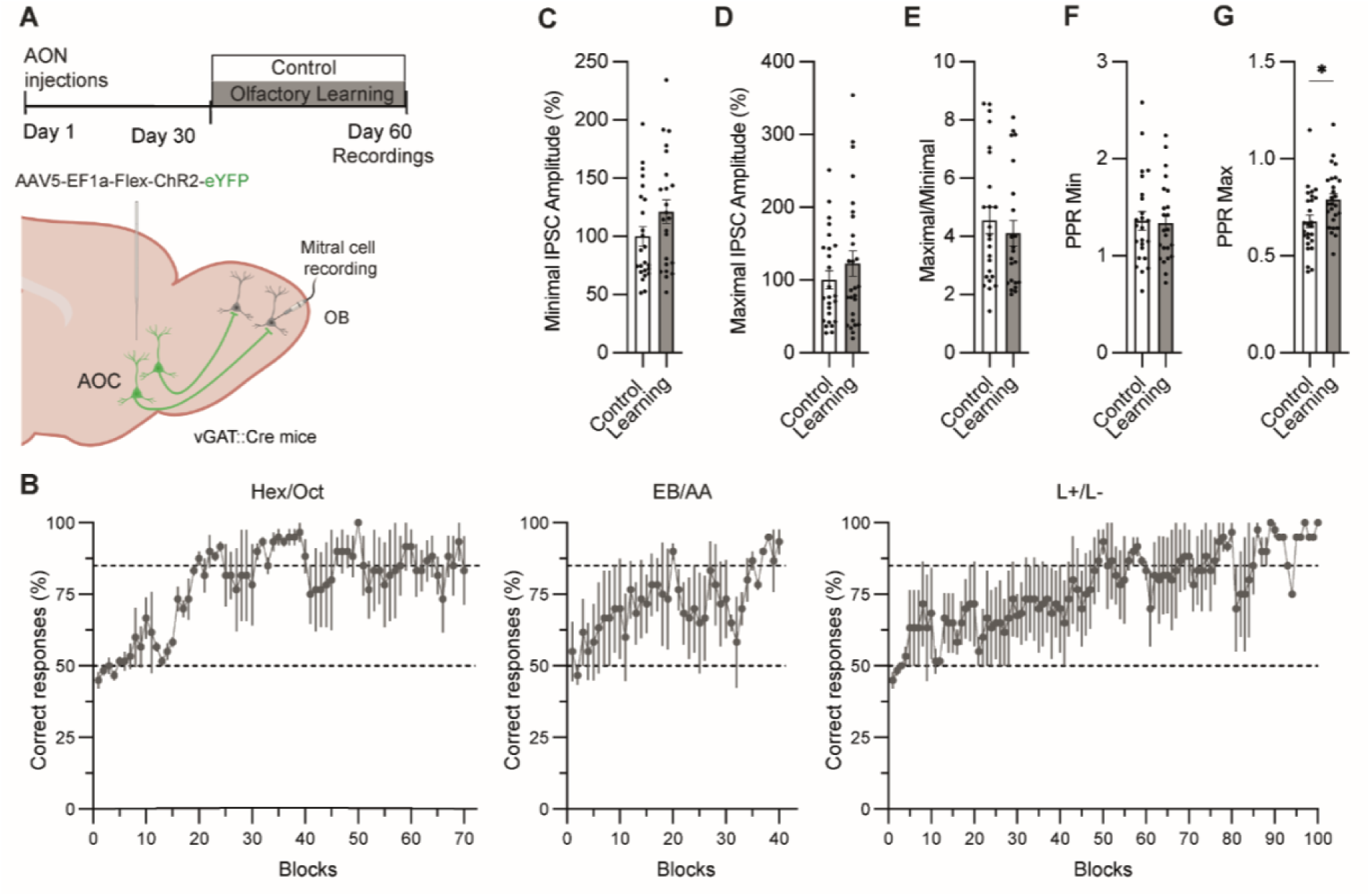
Lack of potentiation of cortical GABAergic inputs onto mitral cells following associative learning. **A**. Upper panel: Chronology of stereotaxic injections and electrophysiology recordings. Lower panel: Stereotaxic injections ChR2-expressing virus in the anterior olfactory cortex (AOC). **B**. Learning performance of three consecutive discrimination tasks (Hexanol vs Octanal; Ethyl Butyrate vs Amyl Acetate; (+)-Limonene vs (-)-Limonene). **C.** Minimal light-evoked IPSCs amplitude (Mann-Whitney test, p=0.146, n=24-23). **D.** Maximal light-evoked IPSCs amplitude (Mann-Whitney test, p=0.478, n=25-27). **E.** Maximal/minimal ratio (Mann-Whitney test, p=0.351, n=24-22) **F**. Paired-pulse ratio for minimal IPSCs (Mann-Whitney test, p=0.903 n=24-23) **G.** Paired-pulse ratio for maximal IPSCs (Mann-Whitney test, p=0.0104, n=26-26). Data is shown as mean ± SEM and individual data points. * p<0.05.

**Supplementary figure 6.**
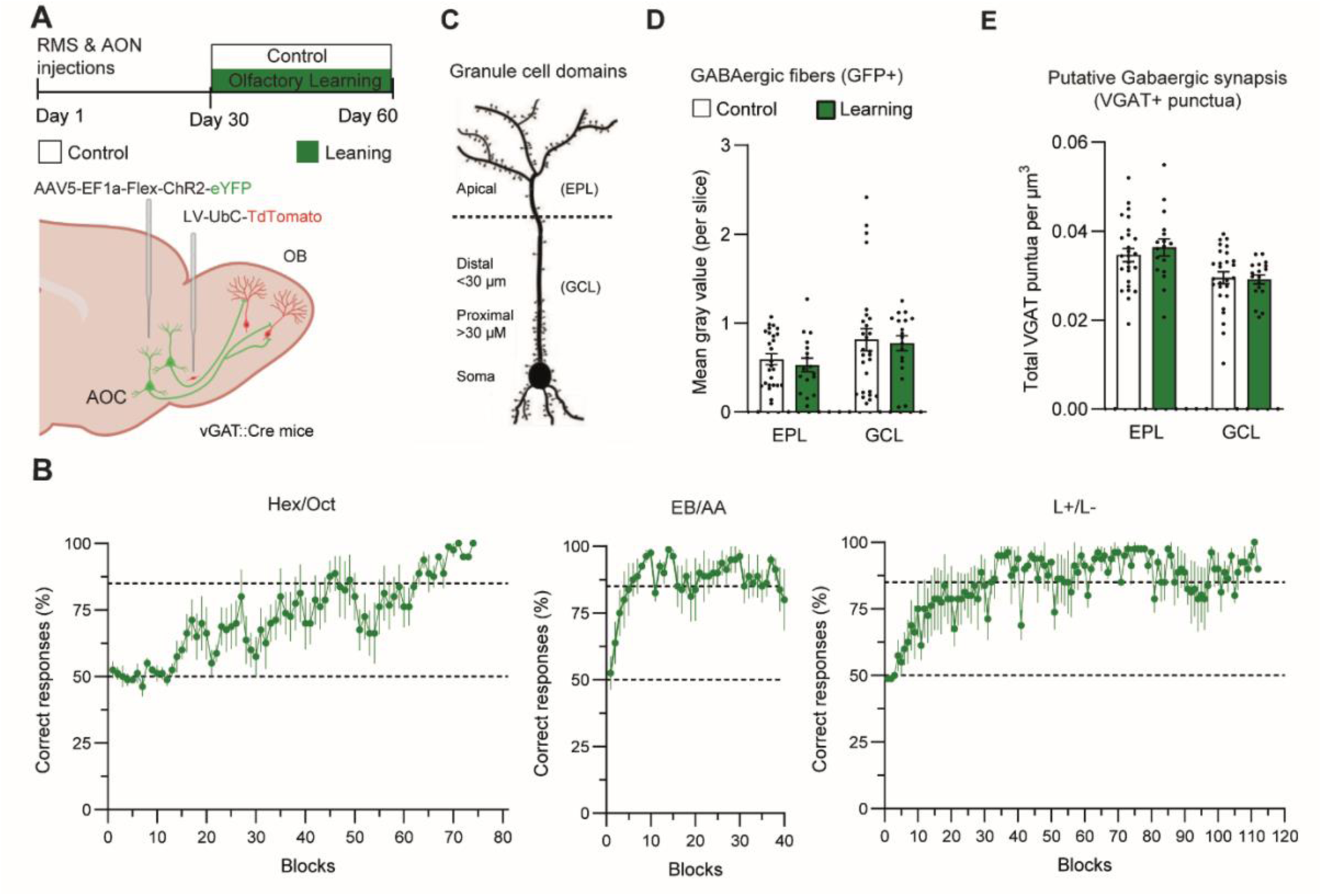
No difference in the total amount of AONp-GABAergic projections and GABAergic putative synapsis after learning in the olfactory bulb. **A**. Upper panel: Timeline of the experiments. Lower panel: stereotaxic injections of TdTomato-expressing virus in the Rostral Migratory Stream (RMS) and ChR2-expressing virus in the Anterior Olfactory Cortex (AOC) in VGAT-CRE mice. **B**. Learning performance of three consecutive discrimination tasks (Hex/Oct = Hexanol vs Octanal; EB/AA = Ethyl Butyrate vs Amyl Acetate; L+/L- = (+)-Limonene vs (-)-Limonene). **C**. Schematic representation of GCs anatomical segmentation and OB layers. **D**. Mean grey value (per OB slice) in both external plexiform layer (EPL) and granule cell layer (GCL) of GABAergic-GFP+ fibers (green channel) in both control and learning group. GABAergic projections signal is reduced in EPL respect to GCL in both groups (Two-way ANOVA, group: F_(1,84)_ = 0.2829 p = 0.5962, layer: F_(1,84)_ = 5.608 p = 0.0202, interaction F_(1,84)_ = 0.0144 p= 0.9048; control EPL vs GCL = p=0.0166; learning EPL vs GCL = p=0.0059, n=17-27 slices from n= 6 mice). **E**. Density of total putative GABAergic synapsis (VGAT+ puncta) in different OB layers. VGAT+ punctua density was higher in EPL respect to GCL in both groups (Two-way ANOVA, group: F_(1,84)_ = 0.2009 p=0.6552, layer: F_(1,84)_ = 16.66 p=0.0001, interaction F_(1,84)_ = 0.4550 p= 0.5018, n=17-27 slices from n= 6 mice). *p<0.05, **p<0.01. Data are shown as mean ± SEM and individual data points.

## Notes

### Competing Interest Statement

The authors have declared no competing interest.

